# Nanosecond methyl dynamics in the eukaryotic RNA exosome core

**DOI:** 10.64898/2026.07.25.740703

**Authors:** Daniela Lazzaretti, Matteo Cagiada, Ayşa Yelboğa, David Stelzig, Kresten Lindorff-Larsen, Till Rudack, Remco Sprangers, Jobst Liebau

## Abstract

Dynamics in proteins occur on a wide range of timescales and are crucial for protein function. On the fast end of that timescale, pico- to nanosecond dynamics have been extensively employed as proxies for entropy and their amplitude can be described by order parameters. Experimentally, NMR can be used to determine order parameters of the protein backbone and, via deuterium relaxation, of methyl groups, yet such experiments cannot be applied to large protein assemblies. In contrast, relaxation-violated coherence transfer experiments, that allow for the determination of side chain order parameters in highly deuterated, methyl-labeled proteins, are more sensitive. Here, we demonstrate that such experiments can be applied to very large, asymmetric protein assemblies by determining axial methyl order parameters for the 300 kDa fully asymmetric core of the eukaryotic RNA exosome complex. Ile-δ1[^13^CH_3_] methyl groups adopt a wide range of order parameters but highly flexible side chains are infrequent. High quality data, which we obtain for flexible regions, is required to observe subtle effects of RNA binding on order parameters. Local cryo-EM Q-scores correlate moderately with order parameters suggesting that Q-scores contain information on nanosecond motions. AF2χ, a recently described prediction tool for side-chain variability, provides good estimates of methyl order parameters, which are, in favorable cases, strongly correlated with experimental values. We thus demonstrate that relaxation-violated coherence transfer experiments can be employed to determine order parameters in large, asymmetric protein complexes that are difficult to capture by other methods, yet are crucial for the understanding of protein function.

**Significance:** Nanosecond side chain dynamics contribute to the entropy of proteins and are therefore proxies for protein stability and binding. Here, we demonstrate that NMR can be employed to experimentally quantify nanosecond dynamics in large, asymmetric proteins paving the way to assess contributions of fast dynamics to the quality of static protein structures. Furthermore, we employ the experimental data to validate computational methods that provide structural insights into nanosecond dynamics.

## Introduction

Dynamics in proteins are crucial for their function, and solution-state NMR is a powerful method to characterize protein motions as it can be employed to study protein dynamics at atomic resolution in solution and on a wide range of timescales, ranging from picoseconds to days (Sheppard et al. 2010; Kay 2011; Kleckner and Foster 2011). On the fast end of that scale, pico- to nanosecond dynamics can be determined from NMR relaxation rates, and their amplitudes represented by order parameters, for which values close to 1 indicate restricted motion and values close to 0 indicate unrestricted motion on that timescale (Lipari and Szabo 1982a, b; Sapienza and Lee 2010; Liao et al. 2012). Backbone order parameters correlate with secondary structure, while there is only weak correlation between methyl side chain order parameters and secondary structure (Sapienza and Lee 2010; Xu and Huang 2021) or solvent accessibility (Mittermaier et al. 2003). Both, side chain and backbone order parameters have been used as proxies to study entropy in proteins (Frederick et al. 2007; Marlow et al. 2010; Sapienza and Lee 2010). Since the entropic component of the free energy is affected by binding events, folding and other structural changes, order parameters are an important measure for thermodynamic properties of proteins (Doig and Sternberg 1995; Yang and Kay 1996; Frederick et al. 2007; Marlow et al. 2010; Kasinath et al. 2013; Sharp et al. 2014; O’Brien et al. 2016). Commonly, backbone order parameters are obtained from ^15^N relaxation experiments, which are, however, limited to relatively small proteins, precluding the study of pico- to nanosecond dynamics in large proteins or protein complexes (Kay 2011; Schütz and Sprangers 2020). ^2^H relaxation experiments have been employed to determine methyl side chain order parameters for proteins as big as ∼80 kDa yet for bigger proteins such experiments are not sufficiently sensitive (Tugarinov et al. 2007).

Sparse methyl labeling approaches in combination with methyl-TROSY NMR experiments (Tugarinov et al. 2003; Kay 2011; Schütz and Sprangers 2020) are substantially more sensitive than ^15^N experiments expanding NMR applications beyond the ∼40 kDa size limit of ‘conventional’ NMR to single-chain proteins bigger than 100 kDa (Overbeck et al. 2022), symmetric protein complexes of up to 1 MDa (Sprangers and Kay 2007) and asymmetric complexes of up to almost half a megadalton (Liebau et al. 2025). Experiments to determine order parameters for such methyl-labeled samples provide the same information as ^2^H relaxation experiments but can be applied to large protein complexes (Tugarinov et al. 2005, 2007; Tugarinov and Kay 2006; Sun et al. 2011). In these relaxation-violated coherence transfer experiments (also termed ‘forbidden’ experiments), double quantum (DQ) or triple quantum (TQ) coherences are generated and their build-up is compared to decay of single quantum (SQ) coherence. The resulting ratio depends on the intra-methyl ^1^H–^1^H cross-correlated relaxation rate, *η*, which in turn can be expressed in terms of the axial methyl order parameter (*S_axis_*^2^, see Eq. 1 and 2 below). The rapidly rotating protons in methyl groups display no homonuclear scalar couplings and in the absence of relaxation *η* = 0, and DQ/TQ coherences cannot be generated (Kay and Prestegard 1987; Tugarinov et al. 2005). However, methyl groups with a large overall correlation time, such as methyl groups in a protein complex, display a fast and a slow relaxation rate, *R*_2 *,H*_*^S^* and *R*_2 *,H*_*^F^* . For such systems 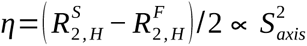 (Tugarinov et al. 2005) facilitating the generation of relaxation-violated (‘forbidden’) DQ/TQ coherences and thus measurement of axial methyl order parameters. Initially, experiments, in which DQ coherences are generated, were developed (Kay and Prestegard 1987; Tugarinov et al. 2005, 2007), followed by the development of TQ-based experiments (Sun et al. 2011), which are more sensitive than DQ experiments and employed in this study.

Side-chain order parameters for methyl-bearing residues are—to a large extent—determined by the residue and methyl type, so that most C_b_ methyl groups in Ala are more rigid than C_g_ methyl groups in Thr, Val and Ile, that in turn are more dynamic than C_d_ methyl groups in Ile and Leu (Best et al. 2004). This is because methyl order parameters depend substantially on the number of rotamers that a side chain samples on a ps–ns timescale, and thus on the number of degrees of freedom between the polypeptide backbone and methyl group (Ming and Brüschweiler 2004; Best et al. 2005). To study the dynamics across a large protein it is thus useful to focus on a single type of methyl group and here we investigate Ile-δ1 methyl groups.

The exosome is a decameric protein complex (Exo10) involved in 3′ to 5′ RNA degradation of linear RNA both in the cytoplasm and in the nucleus (Wasmuth and Lima 2012; Makino et al. 2013; Zinder and Lima 2017). Here, we study the nonameric inactive core, Exo9, of *Chaetomium thermophilum*. The core consists of 6 distinct ring subunits (Rrp41, Rrp42, Rrp43, Rrp45, Rrp46, Mtr3) that form a central pore, which is stabilized by 3 distinct cap subunits (Csl4, Rrp4, Rrp40) that recruit cofactors and substrate RNA (Zinder and Lima 2017), which is then threaded through the central pore and further passed on to the catalytic subunit Rrp44 for degradation (Bonneau et al. 2009; Wasmuth and Lima 2012; Han and van Hoof 2016; Liebau et al. 2025). While it possesses no catalytic activity, Exo9 is essential in eukaryotes (Liu et al. 2006; Lorentzen et al. 2007). Despite its size (Exo9: ∼300 kDa) and full asymmetry, we have previously shown that both methyl and ^19^F NMR can be applied to study interactions, dynamics and structural features of the exosome by employing sparse and modular labeling schemes, in which individual subunits are labeled and reconstituted with NMR-silent subunits (Liebau et al. 2025). This allowed us to characterize functional dynamics of extended loop regions on the millisecond timescale. However, motions on faster timescales remained more challenging to study, although we hypothesized that they are functionally important based on the observation that association of the loop region goes along with increased order parameters as determined from PRE experiments. In addition, measurement of order parameters could report on entropic changes caused by RNA binding.

Structural heterogeneity in single-particle cryo-EM and X-ray crystallography is often interpreted in terms of molecular dynamics, however, typically without reference to timescales of motion. With the advent of time-resolved cryo-EM it has become possible to arrest states with a time resolution of milliseconds (Frank 2017) and, for some approaches, microseconds (Voss et al. 2021; Curtis et al. 2026), allowing to more accurately attribute conformational changes in static structures to timescales of motions. However, to what degree faster motions contribute to structural quality of cryo-EM maps remains largely unexplored and existing approaches rely on computational methods to analyse timescale-unspecific heterogeneity in cryo-EM structures or on MD simulations to propagate motions from static structures (Astore et al. 2025). Since we demonstrate here that ps-ns dynamics can be experimentally determined for structures of the size required for cryo-EM studies, it becomes feasible to site-specifically correlate structural quality factors with order parameters thus allowing to assess whether dynamic data is encoded in static structural data.

Here, we expand the biophysical toolkit for studying large, fully asymmetric protein complexes by demonstrating that axial methyl order parameters are measurable even within these challenging macromolecular complexes. We demonstrate that Ile side chains display a wide range of flexibilities and that high quality data is required to observe subtle effects of RNA substrate binding on pico- to nanosecond Ile side chain dynamics. While it remains challenging to predict methyl order parameters from MD simulations, a recently published method, AF2χ, reliably predicts order parameters when high-quality data is available. Based on our ability to determine methyl order parameters for ‘cryo-EM’-sized complexes, we find a moderate correlation between methyl order parameters and cryo-EM Q-scores suggesting that cryo-EM structural data contains a wealth of information on ps–ns dynamics.

## Results

In order to determine order parameters for Ile-δ1 side chains, spectral assignments are required and, in a first step, we expanded available assignments for the exosome by assigning Ile-δ1 side chains for cap subunit Rrp40. Based on these assignments, we determined and analysed methyl order parameters in the Exo9 complex in the absence and presence of substrate RNA. In order to obtain insights into the structural determinants of order parameters, we next correlated experimental data with computational parameters. Finally, we correlated experimental order parameters with structural quality metrics in order to assess, whether such metrics reflect the presence of pico- to nanosecond motions.

### Ile assignment and RNA interaction of Rrp40

In order to obtain site-specific dynamic information, spectral assignments of methyl groups are required. Previously we assigned methyl-TROSY spectra of Ile-δ1[^13^CH_3_] labeled (I-labeled) Csl4, Rrp41 and Rrp45 reconstituted into Exo9 and observed interactions with RNA (**SI Fig. S1,** (Liebau et al. 2025)). Here, we expanded Ile-δ1 assignments to the cap subunit Rrp40. Resonance assignments for monomeric Rrp40 Ile-δ1 were obtained by point mutations (**Fig. 1a, Table S3B**). Since spectra differ substantially between monomer and Rrp40 in Exo9, Rrp40 point mutants were reconstituted into Exo9 to obtain assignments (**Fig. 1b, Table S3B**). Addition of RNA to Exo9 (**Fig. 1b**) gives rise to only a minor shift for residue I92, which is located in the S1 RNA binding domain of Rrp40, while expected interactions in the KH RNA binding domain are not observable. The chemical shift perturbations caused by RNA binding are notably weaker than for the cap subunit Csl4 (**SI Fig. S1a**) indicating that Ile residues are remote from RNA interaction surfaces or interactions are transient.

**Figure 1:**
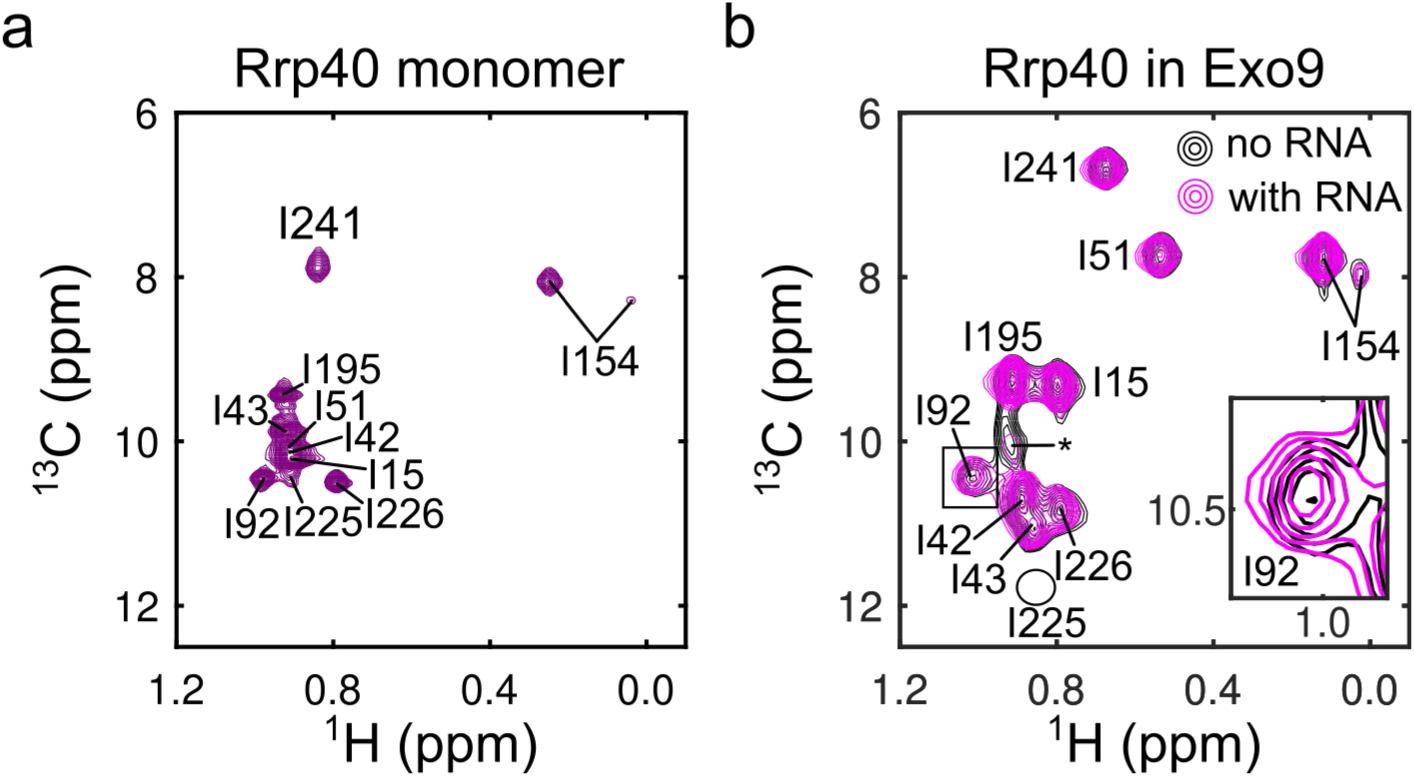
Rrp40 assignment and RNA interaction. **(a)** HMQC of I-labeled monomeric Rrp40. **(b)** HMQC of I-labeled Rrp40 reconstituted into Exo9 in the absence (black) and presence (pink) of RNA. The resonance of Rrp40-I225 is located inside the circle but has very low intensity. * indicates resonances from residual monomeric protein. Inset: Zoom of resonance Rrp40-I92.

### Ile-δ1 display a wide range of nanosecond motions

Spectral assignments in combination with modular, sparse labeling of Exo9 then allowed us to determine methyl order parameters site-specifically. To this end, we individually reconstituted I-labeled Csl4, Rrp40, Rrp41 or Rrp45 into Exo9 and conducted relaxation-violated coherence transfer experiments, for which the build-up of triple-quantum (TQ) coherence relative to the decay of single-quantum (SQ) coherence depends on the axial methyl order parameter *S_axis_*^2^ (**Eq. 1 – 3**).

In total, we determined order parameters for 44 Ile-δ1 methyl groups in Exo9 (**Fig. 2a, b, Fig. S2 and S3, Table S5**) demonstrating that it is indeed possible to determine methyl order parameters for very large, asymmetric protein complexes. However, measurement uncertainties are high, in particular for rigid methyl groups (**Fig. 2b**). The underlying reasons are twofold: 1. The line widths of flexible methyl groups are narrower, giving rise to higher signal-to-noise ratios than for rigid methyl groups. 2. The TQ coherence build-up for rigid methyl groups can be on the same order or faster than the shortest experimental build-up delay (1 ms) implying that the determination of *S_axis_*^2^ becomes unreliable (**Fig. 2c, Fig. S2**). The distribution of order parameters is relatively even, with the exception that very flexible methyl groups (*S_axis_*^2^ < 0.2) are rare (**Fig. 2d**), in line with the general trend of Ile-δ1 methyl order parameters across proteins (Best et al. 2004).

**Figure 2:**
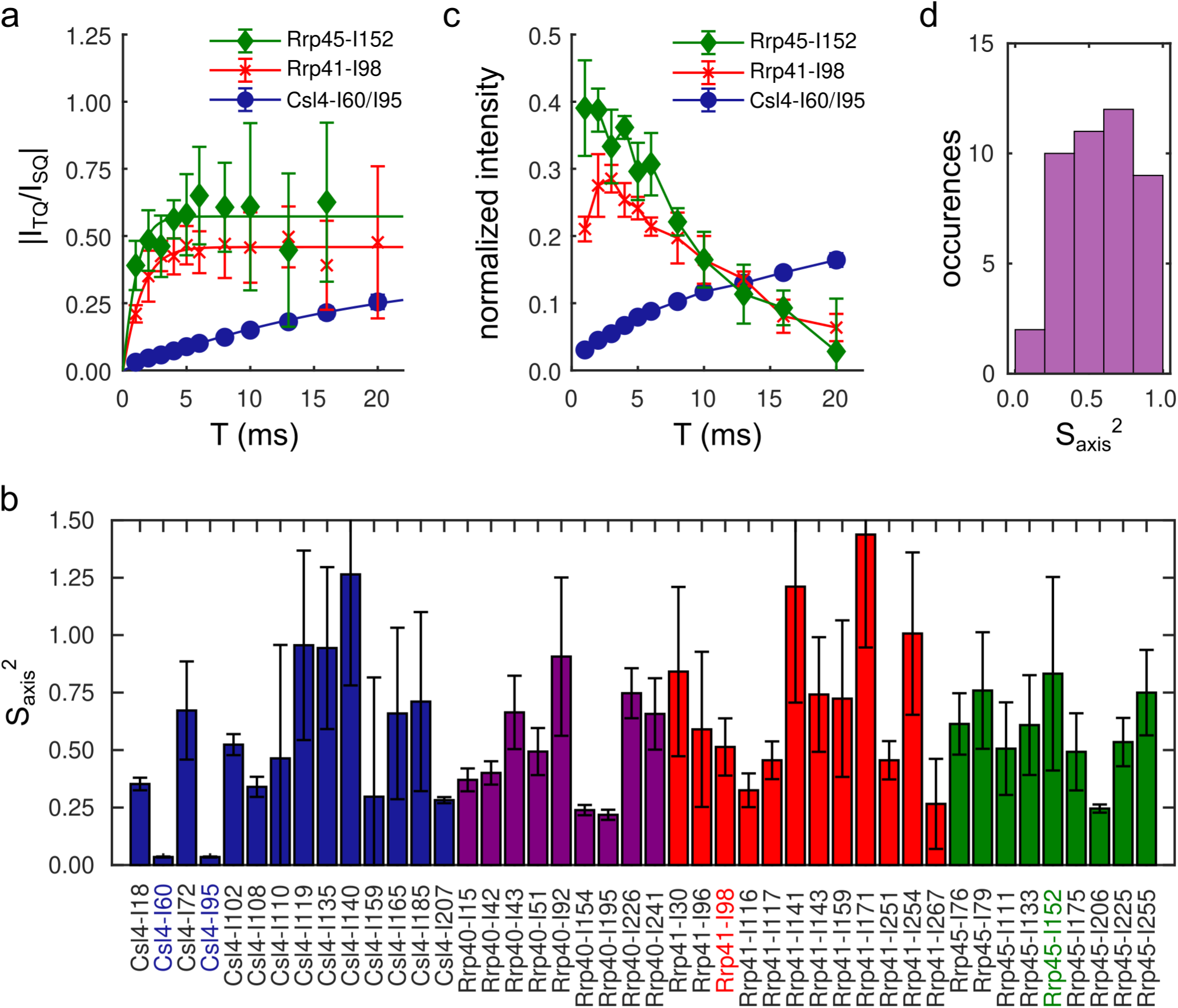
Methyl order parameters for I-labeled subunits in Exo9. **(a)** Representative examples of build-up curves for rigid (Rrp45-I152, green), semi-rigid (Rrp41-I98, red) and flexible (Csl4-I60/I95, blue) Ile residues. The curves are fits using Eq. 1 – 3. **(b)** Axial methyl order parameters determined for Csl4 (blue), Rrp40 (purple), Rrp41 (red) and Rrp45 (green) in Exo9. The residues in **a** and **c** are highlighted. **(c)** Triple quantum coherence build-up curves for Ile residues in **a**. The connecting lines are for guidance. Intensities are referenced to the intensity of the respective SQ coherence (see **Fig. S1**) obtained for *T* = 1 ms. **(d)** Distribution of axial methyl order parameters. Error bars in **a** and **c** are obtained from signal-to-noise ratios of the NMR spectra and represent ±1 SD. Error bars in **b** are obtained from an error analysis as described in the method section and represent ±1 SD.

### Effect of RNA binding on *S_axis_*^2^

Based on axial methyl order parameters determined for apo Exo9, we next sought to test whether RNA substrate binding affects local ps-ns side chain dynamics. We previously demonstrated that Ile-δ1 methyl resonances can be employed to monitor binding of RNA to the exosome via spectral changes (**Fig. S1**) (Liebau et al. 2025). This binding is highly dynamic in the central channel since RNA is not observable there in cryo-EM structures (Makino et al. 2013, 2015; Schuller et al. 2018). To test whether RNA binding also affects ps-ns dynamics, we determined methyl order parameters for the four subunits Csl4, Rrp40, Rrp41 and Rrp45 in Exo9 in the presence of RNA **(Fig. 3, Fig. S4, Fig. S5, Table S5**). Simulation of build-up curves with varying order parameters (**Fig. S6**) show that the initial build-up of |*I*_TQ_/*I*_SQ_| as well as its value for long build-up delays are crucial to distinguish flexible from rigid methyl groups. The latter parameter is difficult to measure, since intensities are lower for longer delay times and thus display higher uncertainties. Build-up curves for flexible side-chains are altered drastically upon changing *S_axis_*^2^, while an identical change of *S_axis_*^2^ for rigid side chains alters build-up curves less drastically. This suggests that changes in *S_axis_*^2^ upon substrate binding are easier to be experimentally observed for flexible side chains, while high quality data is required to observe subtle changes in *S_axis_*^2^ or changes in *S_axis_*^2^ for rigid side chains.

**Figure 3:**
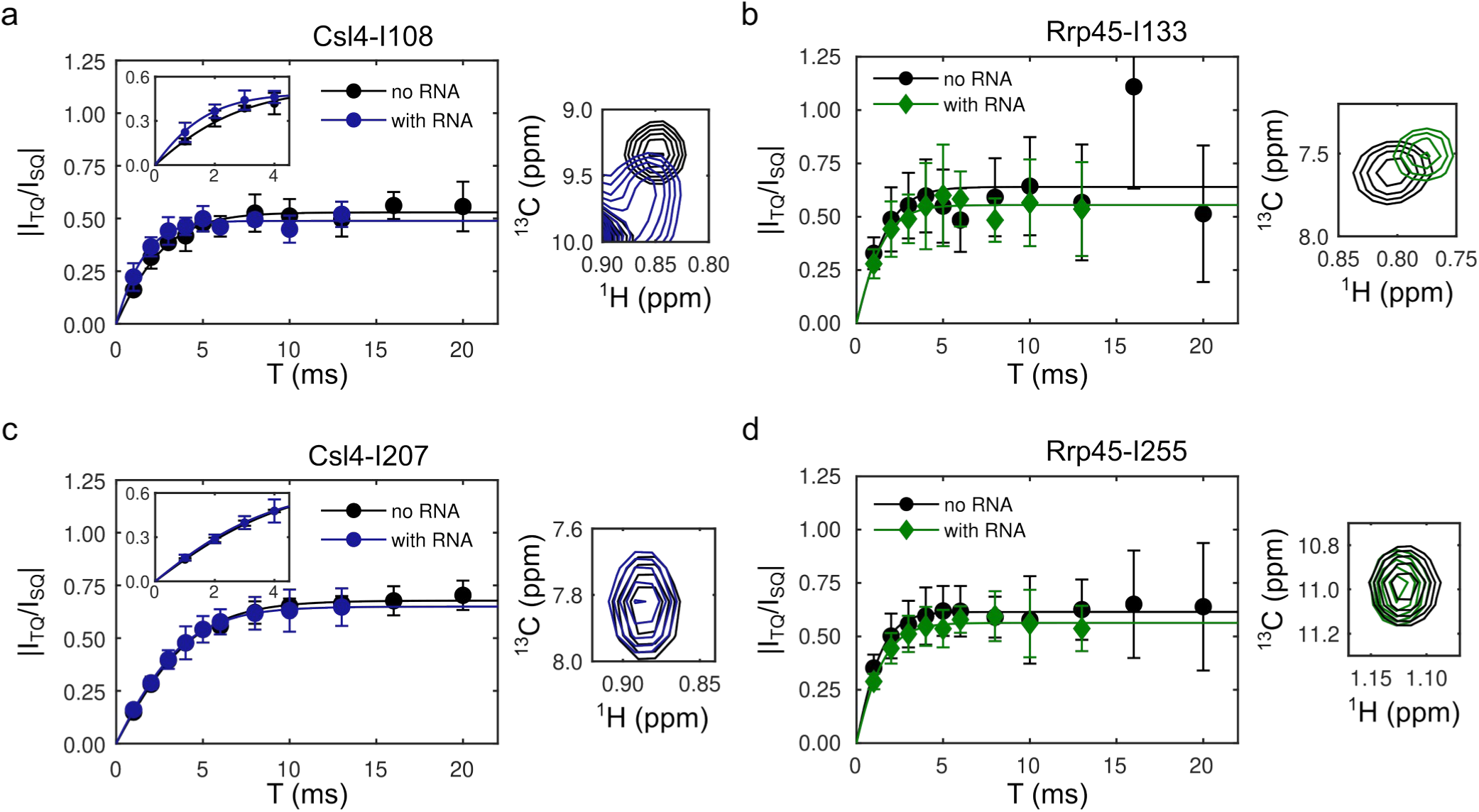
Axial methyl order parameters in the presence of RNA. Representative build-up curves for **(a)** Csl4-I108 and **(b)** Rrp45-I133, which both interact with RNA (inset spectra), and **(b)** Csl4-I207 and **(d)** Rrp45-I255, which both do not interact with RNA (inset spectra), in absence (black) and presence (colored) of a 30mer RNA. The insets in **a** and **c** are a zoom into the initial region of the build-up curves. Error bars are obtained from signal-to-noise ratios of the NMR spectra and represent ±1 SD.

High quality data for the experimental build-up curves can be obtained for rather flexible Ile side chains as shown for Csl4-I108 and Csl4-I207 in **Fig 3a** and **c**. Csl4-I108 probes interactions with RNA (**spectrum in Fig. 3a**) and here we observe a somewhat faster build-up of |*I*_TQ_/*I*_SQ_| in the presence of RNA indicating an increase in rigidity upon RNA interaction (**Fig. 3a inset**). On the other hand, build-up curves are identical for Csl4-I207 (**Fig. 3c inset**) that does not interact with RNA (**spectrum in Fig. 3c**). However, even for Csl4-I108, for which the signal-to-noise ratio is high compared to other Ile-δ1 methyl groups investigated here (compare for example to **Fig. 3b**), an error analysis (see method section) indicates that the determined order parameter is not significantly different in the presence of RNA compared to the apo state. For all other observed Ile side chains build-up curves in the presence and absence of RNA are identical within error limits (**Fig. S5, Table S5**) and, within error limits, *S_axis_*^2^ is unaffected by the addition of RNA, even for residues, which show substantial chemical shift perturbations upon RNA addition (e.g. spectrum in **Fig. 3b**). Thus, RNA binding does not drastically alter *S_axis_*^2^, while high data quality is required to distinguish subtle effects of RNA binding on ps–ns dynamics, which we currently can only achieve in exceptional cases for flexible Ile side chains in large asymmetric complexes. However, as discussed above simulations suggest that it should be feasible to experimentally distinguish more drastic changes in order parameters (**Fig. S6**).

### Structural and dynamic determinants of *S_axis_*^2^

While we thus demonstrated that it is indeed possible to determine *S_axis_*^2^ for large asymmetric complexes, NMR does not provide any direct insights on the nature of the dynamics that are described by *S_axis_*^2^. We, therefore, explored structural and dynamic determinants of axial methyl order parameters on the one hand and how order parameters affect the quality of structural data on the other hand.

We first investigated whether order parameters correlate with the MD-derived solvent-accessible surface area (SASA) of the Ile residues. We observe a moderate negative correlation (ρ = -0.47±0.10, p = 0.0013) between SASA and *S_axis_*^2^ (**Fig. 4a, Table S6**), in line with previous analyses in other proteins (Mittermaier et al. 2003). Of note, |ρ| = 1 is not expected even if the correlation was perfectly monotonic due to measurement uncertainties of *S_axis_*^2^. The maximum correlation expected given the distribution and measurement uncertainties of *S_axis_*^2^, |ρ_ceil_|= 0.72±0.07 for the complete dataset. While surface-exposed Ile residues display lower order parameters than buried ones, buried Ile residues can display a wide range of order parameters between ∼0.5 – 1 (**Fig. S7a**) demonstrating that packing interactions of buried Ile residue do not abolish pico- to nanosecond dynamics. Indeed, this observation is in agreement with the finding that local packing density is an imperfect predictor of order parameters (Ming and Brüschweiler 2004) and with our MD simulations. Qualitatively, Ile-δ1 carbon atoms are not rigid during a 100 ns MD simulation for any of the Ile residues observed here (**Fig. 4b, Fig. S8**). While very flexible residues, such as Csl4-I60, sample a large space, which results in low order parameters, differences in the space sampled between semi-rigid and rigid Ile side chains are qualitatively less obvious (**Fig. 4b**). A quantitative evaluation of the space sampled by Ile-δ1 carbons atoms demonstrates a significant, moderate correlation between RMSF and *S_axis_*^2^ (**Fig. 4a, Table S6, Fig. S7b**). In addition, we calculated axial methyl order parameters from multiple MD simulations and observe a moderate correlation with experimental values (**Fig. S7c, Table S6**) consistent with previous studies (Ming and Brüschweiler 2004; Carbonell and Sol 2009; Gu et al. 2014).

**Figure 4:**
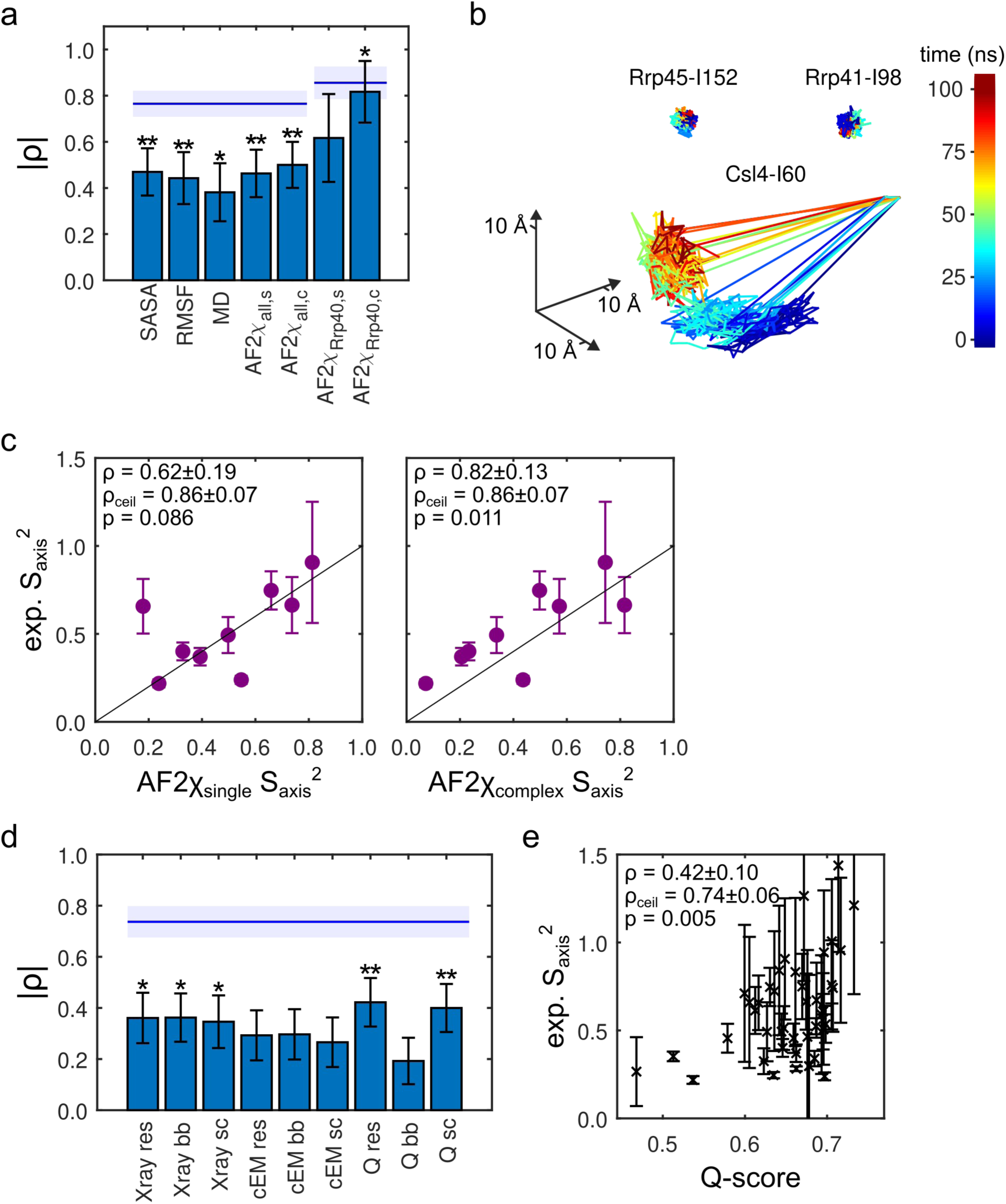
Correlation between axial methyl order parameters and structural parameters. **(a)** Absolute value of Spearman’s rank correlation coefficient ρ for correlations between *S_axis_*^2^ and solvent accessible surface area (SASA) of Ile residues, root-mean-square fluctuations (RMSF) of Ile-δ1 carbon atoms, *S_axis_*^2^ calculated from the MD simulation (MD) and *S_axis_*^2^ predicted by AF2χ using the ‘single’ (s) or ‘complex’ (c) approach (see method section) for all Ile residues and Ile residues of only Rrp40. **(b)** Space sampled by Ile-δ1 carbon atoms from Fig. 1a during a 100 ns MD simulation. Csl4-I60 is very flexible, Rrp41-I98 is semi-rigid and Rrp45-I152 is rigid. **(c)** Correlation between experimental *S_axis_*^2^ of Rrp40 and *S_axis_*^2^ predicted by AF2χ using the ‘single’ (left) and ‘complex’ (right) approach. **(d)** Absolute value of ρ for correlations between *S_axis_*^2^ and various structural parameters (Xray: X-ray resolution, cEM: cryo-EM resolution, Q: Q-score; evaluated for res: full residue, bb: backbone atoms, sc: side chain atoms). **(e)** Correlation between experimental *S_axis_*^2^ and residue Q-scores. Error bars were obtained from error analyses as described in the method section and represent ±1 SD. *p*-values for the correlation between *S_axis_*^2^ and other parameters were evaluated using a two-sided paired-sample *t*-test. **: *p*-value < 0.01, *: 0.01 ≤ *p*-value < 0.05. See **Table S6** for values. The black line in **c** indicates values for which experimental and predicted *S_axis_*^2^ are equal. The blue line (shaded area) in **a** and **d** indicates the maximum expected ρ, ρ_ceil_, (±1 SD), given the measurement uncertainties and distribution of *S_axis_*^2^.

AF2χ has recently been developed as a method to predict conformational side-chain heterogeneity from both predicted and experimentally determined structures, including cryo-EM structures (Cagiada et al. 2025), and has been validated using experimentally determined order parameters and scalar couplings. Briefly, AF2χ uses the internal representations of AlphaFold to identify residues that might populate multiple rotameric conformations, and uses this information to construct an ensemble of configurations to represent the predicted rotamer distributions. Here, we employed AF2χ to predict axial methyl order parameters and compared them to the experimentally determined values. Since it is currently not feasible to use AF2χ to predict *S_axis_*^2^ for the entire Exo9 assembly, we chose two approaches: In the first approach (‘single’), we predicted *S_axis_*^2^ for the isolated subunits. To also account for Ile interactions with neighboring subunits, we, secondly, also predicted *S_axis_*^2^ for sub-assemblies, where ensembles were generated for a target subunit and all its direct interaction partners (‘complex’ approach). Overall, we observe a significant but moderate correlation between predicted and calculated *S_axis_*^2^, which is somewhat improved if neighbouring subunits are taken into consideration (**Fig. 4a, SI Fig. S9a)**. A reason for the moderate correlation might be the high experimental uncertainty, as well as limitations in AF2χ that include a lack of incorporation of timescale information in the comparison and are discussed in detail elsewhere (Cagiada et al. 2025). Since we noticed substantial differences in performance of AF2χ for different subunits, we also analysed subunits individually. For Rrp40 we observe a strong correlation between experimental and predicted *S_axis_*^2^ using the ‘complex’ approach **(Fig. 4c)**. For the ring subunits, Rrp41 and Rrp45, the correlation is weaker, while we observe almost no correlation for Csl4 **(SI Fig. S9b-e)**. The reason for these differences in correlation is likely rooted in the fact that the ring subunits contain mostly rigid Ile side chains, for which the experimental uncertainties of *S_axis_*^2^ are high and for which accurate predictions are more challenging. For Csl4 experimental measurement uncertainties are overall high (mean *S_axis_*^2^ SD 0.24), while for Rrp40 the experimental data quality is best of all subunits (mean *S_axis_*^2^ SD 0.11). Subunit-specific variations in data quality are also reflected by relatively low ρ_ceil_ for Csl4 and Rrp45, which suggests that due to high measurement uncertainties and close clustering of *S_axis_*^2^ a high correlation can not be expected. For Rrp40 the correlation is improved if neighboring subunits are taken into consideration, suggesting that the interaction surface affects axial methyl order parameters. For this approach, the prediction by AF2χ outperforms other computational approaches of predicting of *S_axis_*^2^ and is, indeed, close to ρ, the maximum correlation expected.

### Cryo-EM Q-scores correlate with *S_axis_*^2^

While it remains challenging to identify structural determinants of methyl order parameters, we hypothesized that static structural data might directly contain information on nanosecond dynamics. Since we here demonstrate that methyl order parameters can be determined for protein assemblies, for which cryo-EM structures are available, we sought to determine whether ps–ns motions are correlated with loss of structural quality for both X-ray and cryo-Em structures of ctExo9. We tested the correlation between *S_axis_*^2^ and structural metrics by correlating X-ray B-factors, cryo-EM local resolution and cryo-EM Q-scores each evaluated for the entire Ile residue, the Ile backbone or the Ile side chain, with axial methyl order parameters. B-factors are a measure of mean atom displacement in X-ray crystallography and here we obtained B-factors from our recently published X-ray structure of Exo9 (PDB: 8PEL, (Liebau et al. 2025)). The cryo-EM Q-score describes the agreement between cryo-EM electron densities and a modeled map. Recent studies further demonstrate that, among other factors, local flexibilities contribute to the Q-score (Pintilie et al. 2020, 2025). Cryo-EM Q-scores and local resolution were obtained from our recently published cryo-EM structure of Exo9 (PDB: 8R1O, (Liebau et al. 2025)). We find that order parameters and cryo-EM Q-scores when evaluated for the entire residue show the strongest correlation of all parameters tested here (*ρ* = 0.42±0.10, p = 0.005) (**Fig. 4d+e, Fig. S10, Table S6**) yet correlations are moderate, suggesting major contributions from other effects to the Q-scores. Conversely, cryo-EM local resolution is not a good predictor for methyl order parameters (p-values > 0.05), while X-ray backbone B-factors show correlations that are weaker than for the residue Q-score (**Fig. S10, Table S6**).

## Discussion

Order parameters are an important proxy for entropy in proteins (Kasinath et al. 2013) and are thus an important parameter for the investigation of protein folding and stability and they can be employed to study molecular interactions (Frederick et al. 2007) and side chain packing (Ming and Brüschweiler 2004; Carbonell and Sol 2009; Sharp et al. 2014). Their determination has therefore been of interest since seminal contributions (Halle and Wennerström 1981; Lipari and Szabo 1982a, b; Lipari et al. 1982). Methodological advances that rely on the measurement of relaxation-violated, double/triple-quantum coherence build-up have made it possible to study methyl order parameters in large protein complexes (Sun et al. 2011). Nonetheless, reports are limited to small proteins, model proteins or large symmetric complexes (Tugarinov et al. 2005, 2007; Sun et al. 2011; Capdevila et al. 2018; Wang et al. 2019; Dubey et al. 2021; Xu and Huang 2021). Here, we demonstrate that relaxation-violated coherence transfer experiments in combination with modular methyl-labeling facilitate the determination of methyl order parameters in very large, fully asymmetric protein complexes. We find that Ile-δ1 methyl order parameters are broadly distributed with the exception that highly flexible methyl groups are rare. We further observe that RMSF, SASA and cryo-EM residue Q-score are moderately correlated with order parameters. Predicted order parameters are also in moderate agreement with experimental data. Our approach is limited due to a currently small dataset and in that measurement uncertainties, in particular for rigid methyl groups, are relatively large owing to limitations in spectral quality and fast build-up of triple-quantum coherences.

Previous reports establish that methyl order parameters can be employed to study binding between protein and substrate (Frederick et al. 2007). Our data suggest that in large systems substrate binding can be observed in favorable cases, or possibly when changes in order parameter are large. However, due to large measurement uncertainties, subtle changes in order parameters are beyond the detection limit for very large, asymmetric protein complexes and consequently we cannot observe effects of RNA binding except in one favorable case.

To date it remains challenging to identify structural and dynamic determinants of axial methyl order parameters, precluding accurate predictions, and correlations between experimental data and theoretical predictions are typically moderate (Ming and Brüschweiler 2004; Carbonell and Sol 2009; Gu et al. 2014). It has previously been acknowledged that computational methyl order parameters depend on the force field (O’Brien et al. 2016) and a number of approaches have been developed that improve on the standard method for order parameter calculation from MD simulations and reach better predictive power (Hoffmann et al. 2018a, b), or alternatively use MD simulations to interpret experimentally measured relaxation rates without resorting to model fitting (Kümmerer et al. 2021). Inaccuracies might arise from inadequate parametrization (Hoffmann et al. 2018a), insufficient sampling (O’Brien et al. 2016; Hoffmann et al. 2018a) and complex dynamic behavior, such as anisotropic protein reorientation (Hoffmann et al. 2018b) or multiple correlation times (Zumpfe et al. 2024) resulting in deviations from experimental values. Further, a more direct comparison between experiments and simulations would require direct calculations of the measured relaxation rates, thus also taking into account variation in motional timescales and the timescale of overall tumbling (Gu et al. 2014; Lindorff-Larsen et al. 2016; Hoffmann et al. 2018a, b). Finally, experimental order parameters are derived using motional models that may not fully capture the complex motions in a protein.

It would be desirable to facilitate order parameter estimates based on easy-to-access structural parameters. Efforts have been undertaken to achieve this, among others by implementing methyl order parameter prediction directly from static three-dimensional structures (Ming and Brüschweiler 2004; Carbonell and Sol 2009; Cagiada et al. 2025). Here, we show that measures such as SASA or AF2χ have moderate predictive power and are in particular valuable to distinguish flexible from rigid methyl groups. If MD simulations are available, RMSF and direct calculation of order parameters provides a moderate correlation with experimental order parameters. In some favorable cases, as for Rrp40, AF2χ provides predictions that are in very good agreement with experimental data. However, while predictions seem to be better for dynamic regions and when experimental uncertainties are low, it remains difficult to identify criteria that allow for an *a priori* decision on whether order parameters can be reliably predicted for a protein. It seems likely that prediction methods will improve with an increased experimental data foundation.

Investigations of contributions of ps–ns dynamics to structural heterogeneity in cryo-EM structures remain scarce. Order parameter determination by NMR would, in principle, facilitate such studies, however, almost all cryo-EM structures deposited in the PDB are bigger than ∼60 kDa and ∼90% of all cryo-EM structures are >100kDa and thus substantially bigger than the size limit for conventional NMR. We overcome this limitation by employing modular and sparse methyl labeling of Exo9, allowing us to determine axial methyl order parameters in the 300 kDa asymmetric complex. Together with the high-resolution cryo-EM structure of Exo9, experimental determination of *S_axis_*^2^ by NMR thus facilitates the correlation of metrics of structural heterogeneity with experimental axial methyl order parameters. The correlation we here observe between Q-score and order parameters suggests that nanosecond dynamics are indeed contributing to structural heterogeneity, however the correlation is only moderate. As previously reported, factors that contribute to reduced Q-scores, aside from molecular motions, are conformational heterogeneity from motions slower than nanoseconds, radiation damage and in particular poor model fitting (Pintilie et al. 2020, 2025). With our limited dataset we cannot currently deconvolute such effects from molecular motions. Nonetheless, this study provides an experimental indication that nanosecond dynamics are encoded in structural quality metrics, suggesting that a treasure of dynamic information lies hidden in structural data. To lift this treasure, further efforts will be required to deconvolute order parameters from other contributions to structural quality metrics.

## Conclusions

The role of pico- to nanosecond dynamics for the function of enzymes has long been acknowledged in that such motions contribute to the entropic component of the free energy, which is altered by and thus a determinant of functionally relevant events such as substrate binding, protein-protein interactions or folding/unfolding transitions. Quantification of such fast dynamics is thus of key importance for a molecular understanding of protein function and here we demonstrated that such dynamics can be determined in very large, asymmetric complexes and correlate with structural factors and cryo-EM data quality metrics suggesting that cryo-EM datasets contain information on ps–ns timescale dynamics. However, disentangling dynamic contributions from other contributions to cryo-EM quality metrics remains challenging. Other challenges remain in understanding the nature of molecular motions that determine methyl order parameters. Expansion of datasets of experimentally determined axial methyl order parameters for proteins with known cryo-EM structures in combination with MD simulations will prove valuable in advancing those insights.

## Materials and Methods

### Expression, methyl labeling and purification

*Chaetomium thermophilum* exosome subunits (**Table S1**) were prepared as described previously (Liebau et al. 2025). Briefly, subunits were expressed individually or, for Rrp43 and Rrp46, in pairs, in D_2_O M9 medium employing recycled D_2_O and ^1^H^12^C glucose as carbon source for non-labeled subunits. Ile-δ1[^13^CH_3_]-labeled (I-labeled) subunits were expressed in fresh D_2_O M9 medium employing ^2^H^12^C glucose and 60 mg/L ^2^H^12^C ketobutyric acid-4-^13^CH_3_, which was added 1 h prior to induction with isopropyl-β-D-thiogalactopyranosid (IPTG). Protein was expressed overnight at 20 °C, cells were harvested, resuspended in 50 mM sodium phosphate buffer, pH 7.4, 150 mM NaCl, 10 mM imidazole (buffer A), lysed by addition of Tri ton X100 and lysozyme and by sonication and debris was removed by centrifugation. The supernatant was applied onto a Ni-NTA column and protein was eluted in buffer A supplemented with 300 mM imidazole. 1 mg of TEV protease was added to the sample, which was dialysed overnight into 20 mM HEPES, pH 7.5, 150 mM NaCl (buffer B). TEV protease and the cleaved His-tag were removed by an inverse Ni-NTA column and the protein was subjected to size-exclusion chromatography into 10 mM HEPES buffer pH 7.5, 200 mM NaCl (buffer C).

### RNA *in vitro* transcription

30mer linear RNA (**Table S2**) was prepared by *in vitro* transcription and purified by anion exchange chromatography using a preparative DNAPac 100 column (Dionex) as described previ ously (Liebau et al. 2025).

### NMR sample preparation

To reconstitute Exo9, the individual subunits were pooled in stoichiometric amounts, incubated for 30 minutes and subjected to size-exclusion chromatography. The fractions containing the exosome complex were pooled and concentrated. Next, the sample was diluted 75x in D_2_O-based buffer C. For RNA-containing samples, RNA was added in 1.5x excess over the exosome complex and the sample was incubated for 30 minutes. Next, the sample was concentrated and the dilution-concentration procedure was repeated once to obtain the final NMR sample.

Experiments were conducted in NMR tubes with a diameter of 3 mm or in 5 mm Shigemi tubes at 313 K. The final NMR sample volume was ∼180 µl (3 mm tubes) or ∼260 µl (5 mm Shigemi tubes) containing 100–200 µM Exo9, in which one subunit (Csl4, Rrp40, Rrp41 or Rrp45) was I-labeled, in D_2_O-based buffer C with 1.5x excess 30mer RNA where applicable.

### Rrp40 Ile-δ1 assignments

Assignment mutants for Rrp40 were prepared by individually mutating each of its 10 Ile residues into Val residues using a Quikchange-like procedure. Primers are listed in **Table S3A**. Subsequently, proteins were over-expressed, purified and reconstituted as described above and a methyl-TROSY HMQC was obtained for the monomeric mutant and the mutant reconstituted into Exo9 to obtain assignments as described previously (Liebau et al. 2025).

### NMR experiments

Monomer assignment experiments were conducted on a Bruker 600 MHz Avance Neo spectrometer (14.1 T magnetic field strength) equipped with a nitrogen-cooled triple resonance TCI probehead. 2D methyl-TROSY experiments were acquired employing the SOFAST-HMQC experiment (Amero et al. 2009), with 1024 x 400 complex points, a carbon acquisition time of 66 ms and an interscan delay of 0.5 s. All other NMR experiments were conducted on a Bruker 800 MHz Avance Neo spectrometer (18.8 T magnetic field strength) equipped with a helium-cooled triple resonance TCI probehead. 2D methyl-TROSY experiments were acquired employing the SOFAST-HMQC experiment, with 1024 x 96 complex points, a carbon acquisition time of 9.2 ms and an interscan delay of 0.5 s. Relaxation-violated coherence transfer experiments were conducted using the pulse sequence developed by Sun et. al (Sun et al. 2011) implemented in the Bruker pulse sequence library as ‘memqtqhmqc3d’, for which the intensity of the SQ and TQ coherence, *I*_SQ_ and *I*_TQ_, respectively, present after a variable build-up delay *T* (*T* = [1, 2, 3, 4, 5, 6, 8, 10, 13, 16, 20] ms without RNA; *T* = [1, 2, 3, 4, 5, 6, 8, 10, 13] ms with RNA) was measured in an interleaved manner. Experiments were acquired with a carbon acquisition time of 9.2 ms, an interscan delay of 1.2 s and 96 transients were averaged. **Table S4** lists samples and NMR experiments conducted on them.

NMR data were processed using the NMRPipe/NMRDraw software suite (Delaglio et al. 1995). Noise was estimated using NMRPipe’s specStat script. Measurement uncertainties for the intensities *I*_SQ_ and *I*_TQ_ were obtained from at least 3 experimental repeats. Intensities were retained for further analysis if they were at least 1.5x above noise level.

Axial methyl order parameters *S*_axis_ ^2^ were obtained as previously described (Tugarinov et al. 2007; Sun et al. 2011):

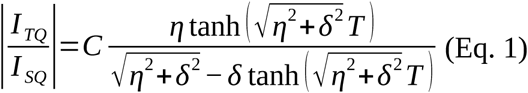

for which *C* = ¾ for TQ-based experiments and *η* is the intra-methyl ^1^H–^1^H cross-correlated relaxation rate given by:

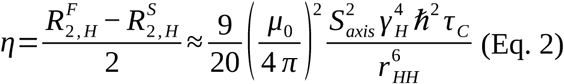

in which 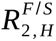 are the fast and slow relaxation rate of the methyl proton single quantum transitions, *μ*_0_ is the vacuum magnetic permeability, *γ*_H_ is the gyromagnetic ratio of protons, *τ*_C_ is the rotational correlation time of the complex, *ħ* is the reduced Planck constant and *r*_HH_ is the distance between pairs of protons (1.813 Å). *δ* is a parameter accounting for relaxation due to external protons (Tugarinov et al. 2007):

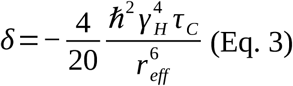

for which the distance between the methyl group and all external protons is represented by a single proton located at a distance *r*_eff_.

As evident from Eq. 2, *η* depends on both *S_axis_*^2^ and *τ*_C_ implying that absolute values for *S_axis_*^2^ can only be obtained if *τ*_C_ is known. However, in this study, we do not compare order parameters between differently-sized complexes such that *τ*_C_ can be treated as a constant scaling factor, which we assumed to be 200 ns giving rise to order parameter values between 0 – 1 within error limits.

### Molecular Dynamics Simulation

To initiate the molecular dynamics (MD) simulations, we utilized complete structural models of the *C. thermophilum* Exo9 complex in both the open and closed Rrp42-EL states. These models (available at https://doi.org/10.5283/EPUB.77450) were generated and prepared as simulation system following previously established protocols(Liebau et al. 2025). Briefly, the structures were constructed by merging the experimental cryo-EM structure of the Exo9 complex (PDB ID: 8R1O) with AlphaFold2-based predictions to model the missing loop regions.

Simulation systems were prepared as previously described (Liebau et al. 2025). MD simulations were performed using the OPLS/all-atom force field (Jorgensen et al. 1996) within GRO-MACS 2021 (Abraham et al. 2015) in aqueous environment. The systems were gradually heated to room temperature (293.15 K) over 1 ns in a nVT ensemble, using a 1 fs time step size, with the temperature maintained constant via the V-rescale thermostat (Bussi et al. 2007). Equilibration continued with a 1 ns nVT run at 293.15 K, followed by a 10 ns nPT run using a Berendsen barostat (Berendsen et al. 1984) and V-rescale thermostat (Bussi et al. 2007). Subsequently, a 100 ns NPT productive run with step size of 2 fs was performed for each system us ing a Nosé-Hoover thermostat (Nosé 1984; Hoover 1985) and Parrinello-Rahman barostat (Parrinello and Rahman 1981). Detailed simulation parameters followed established protocols (Liebau et al. 2025). For each conformation, three independent simulation replicates were carried out in GROMACS 2021 (Abraham et al. 2015), resulting in a total of six trajectories of 100 ns each. All mdp parameter files are available at https://doi.org/10.5283/EPUB.77450.

### Simulation evaluation

To assess the structural stability of the simulations, standard evaluation metrics were applied, including analysis of the backbone root-mean-square deviation (RMSD) of the Cα atoms across the trajectory. Further conformational changes, changes in the interaction network, and the secondary structure were monitored using kontakteUR (Scherlo et al. 2026). These analyses were used to ensure all trajectories reached stable conformational ensembles suitable for subsequent evaluation of side chain dynamics.

To analyze isoleucine side-chain motions, each 100 ns trajectory was first fitted to the protein backbone of the first frame of the respective trajectory using `gmx trjconv -fit rot+trans`, thereby removing global translational and rotational motions. Only the equilibrated second half of each production run (50–100 ns) was used for quantitative analysis to ensure equilibration.

Root-mean-square fluctuations (RMSF) of isoleucine side chains were computed using GROMACS 2021 (`gmx rmsf`). Multiple atom selections were tested, including all non-hydrogen side-chain atoms, all side-chain atoms, methyl-group atoms of the isoleucines of interest and all atoms of each residue. RMSF of methyl groups provided the best correlation with experimental data and are reported here.

Solvent-accessible surface areas (SASA) of selected isoleucine side chains were calculated using the `gmx sasa` module of GROMACS 2021 (Bondi 1964; Eisenhaber et al. 1995). A probe radius of 0.14 nm, corresponding to a water molecule, was used throughout. The SASA was calculated for the whole complex, and averaged for each residue. Values for isoleucine residues of interest were selected subsequently.

To obtain statistically robust estimates, SASA values were computed independently for six simulation replicates. For each replicate, `gmx sasa` outputs a time series from which the program internally calculates a mean and standard deviation over the trajectory. The reported ensemble mean was computed as the arithmetic mean across all six per-replicate means. The over all standard deviation was derived by pooling the six individual standard deviations according to

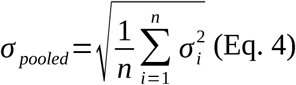

where *n*=6 and σ_i_ is the standard deviation reported by GROMACS for each replicate.

### Calculation of methyl order parameters from predicted structural ensembles

Predicted methyl order parameters were computed from structural ensembles generated using AF2χ (Cagiada et al. 2025), starting from the cryo-EM structure of *C. thermophilum* Exo9 (PDB: 8R1O). Predictions were performed independently for each of the four subunits of interest (Csl4, Rrp40, Rrp41, and Rrp45) under two distinct structural contexts: (i) the subunit in isolation (AF2χ_single_ *S*_axis_^2^), and (ii) the subunit in complex with its neighbouring subunits within the exosome complex (AF2χ_complex_ *S*_axis_^2^). Neighbouring subunits were defined as any protein chain in the complex, for which more than 10% of residues were in contact with the target subunit (C^α^–C^α^ distance < 7 Å).

Structural ensembles were generated using AF2χ with the following settings: a custom template derived from the cryo-EM structure (no multiple sequence alignment and template-only mode), and predictions drawn exclusively from AF2 models 1 and 2. Each ensemble consisted of 50 predicted structures.

Methyl order parameters were then calculated from these side-chain coordinate ensembles using the following equation (Henry and Szabo 1985):

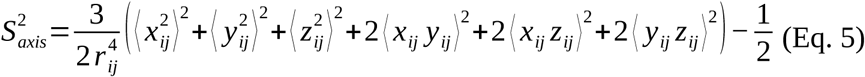

where the components of the interatomic bond vector between carbon *i* (bonded to a methyl group) and carbon *j* (in a methyl group) are denoted by *x*_ij_, *y*_ij_ and *z*_ij_; The length of the C–C bond is denoted by *r*_ij_ and the angle brackets denote averaging over the ensemble structures (Henry and Szabo 1985).

### Calculation of methyl order parameters from MD simulation

Methyl order parameters were computed from the same MD simulations as RMSF and SASA calculations, for which all frames were aligned to the C_α_ atoms to remove global tumbling. For the calculation, only the second half of each trajectory was used to ensure equilibration of the system,employing Eq. 5, for which *x*_ij_, *y*_ij_ and *z*_ij_ denote the coordinates of a vector of unit length (*r*_ij_ = 1) connecting the Ile-δ1 carbon atom and the center of mass of the methyl protons and the time average was computed over the 50 ns trajectory in steps of 5 ps. Of note, alternative methods that achieve better correlation to experimental order parameters have been developed, such as methods that block-average over varying intervals of the MD simulation (Markwick et al. 2007), include overall molecular tumbling (Kümmerer et al. 2021), employ optimized force fields (Hoffmann et al. 2018a) or directly calculate relaxation rates (Hoffmann et al. 2018b, 2020; Kümmerer et al. 2021). Implementation of such approaches is beyond the scope of this experimental study and we thus do not explore them here.

### Correlation statistics

Correlation between *S*_axis_^2^ and other parameters was assessed using Spearman’s ranked correlation coefficient *ρ* to account for possible non-linearity in the correlation. The *p*-value was evaluated using a two-sided paired-sample *t*-test. The maximum expected ρ, given the measurement uncertainties and distribution of *S_axis_*^2^, ρ_ceil_, was determined by assuming a perfectly monotonic relationship between experimentally determined *S_axis_*^2^ and the correlated parameter. 1000 artificial datasets were generated within the measurement uncertainty of *S_axis_*^2^, conserving the experimental distribution of *S_axis_*^2^ values, and Spearman’s ranked correlation coefficient was determined for each artificial dataset. ρ_ceil_ was estimated from the mean and the uncertainty from the standard deviation.

### Experimental uncertainties

Experimental uncertainties for *S*_axis_^2^, *r*_eff_ and *ρ* were obtained by creating 1000 artificial data sets within the standard deviation of the experimental intensities (for *S*_axis_^2^ and *r*_eff_) or *S*_axis_^2^ (for *ρ*) using a Gaussian weighting function and analysis of the data as described above. The standard de viations for the parameters of the artificial data represent the experimental uncertainty.

## Acknowledgments

We thank Nadine Stefan for excellent technical assistance and Torben Fürtges for support with the analysis of the MD simulation trajectories. Jan Overbeck and Philip Wurm (Bruker Biospin) are acknowledged for support in conducting NMR experiments. All present and past group members are acknowledged for critically discussing the results during the course of the project.

## Funding

This project is funded from the European Union’s Horizon 2020 research and innovation programme under the Marie Skłodowska-Curie grant agreement No. 89550 (to JL), by the German Research Foundation (Deutsche Forschungsgemeinschaft) under grant agreement No. SP 1324/3-1, by the European Research Council under the European Union’s Seventh Framework Programme (FP7/2007–2013), ERC grant agreement No. 616052 (to RS) and by a Novo Nordisk Foundation Postdoctoral Fellowship (NNF23OC0082912; to MC). We acknowledge access to computational resources via a grant from the Carlsberg Foundation (CF21-0392; to KLL).

## Authors contributions

Conceptualization: RS, JL, DL

Data curation: JL, MC, AY

Formal Analysis: JL, DL, AY, MC

Funding acquisition: RS, JL, TR, MC, KLL

Investigation: JL, DL, DS, AY, TF, TR, MC

Methodology: RS, JL, DL, MC, KLL

Project administration: RS, JL, TR, KLL

Supervision: RS, JL, TR

Visualization: JL

Writing – original draft: RS, JL, DL, TR, AY, MC, KLL

Writing – review & editing: RS, JL, DL

## Competing interests

The authors declare no competing interests.

## Data and materials availability

All data are available in the manuscript or the supplementary materials. These structural models generated with MD simulations are publicly accessible at https://doi.org/10.5283/EPUB.77450. The structural ensemble generated with AF2χ, code and script used to calculate methyl order parameters are available at: https://github.com/matteo-cagiada/_2026_Lazzaretti_Nanosecond_.

## Notes

### Competing Interest Statement

The authors have declared no competing interest.

## References

1. Abraham MJ, Murtola T, Schulz R, et al (2015) GROMACS: High performance molecular simulations through multi-level parallelism from laptops to supercomputers. SoftwareX 1–2:19–25. 10.1016/j.softx.2015.06.001

2. Amero C, Schanda P, Durá MA, et al (2009) Fast two-dimensional NMR spectroscopy of high molecular weight protein assemblies. J Am Chem Soc 131:3448–3449. 10.1021/ja809880p

3. Astore MA, Silva-Sánchez D, Blackwell R, et al (2025) Comparing cryo-EM methods and molecular dynamics simulation to investigate heterogeneity in ligand-bound TRPV1. Nat Commun 17:1067. 10.1038/s41467-025-67821-2

4. Berendsen HJC, Postma JPM, van Gunsteren WF, et al (1984) Molecular dynamics with coupling to an external bath. J Chem Phys 81:3684–3690. 10.1063/1.448118

5. Best RB, Clarke J, Karplus M (2004) The Origin of Protein Sidechain Order Parameter Distributions. J Am Chem Soc 126:7734–7735. 10.1021/ja049078w

6. Best RB, Clarke J, Karplus M (2005) What Contributions to Protein Side-chain Dynamics are Probed by NMR Experiments? A Molecular Dynamics Simulation Analysis. Journal of Molecular Biology 349:185–203. 10.1016/j.jmb.2005.03.001

7. Bondi A (1964) van der Waals Volumes and Radii. J Phys Chem 68:441–451. 10.1021/j100785a001

8. Bonneau F, Basquin J, Ebert J, et al (2009) The yeast exosome functions as a macromolecular cage to channel RNA substrates for degradation. Cell 139:547–559. 10.1016/j.cell.2009.08.042

9. Bussi G, Donadio D, Parrinello M (2007) Canonical sampling through velocity rescaling. J Chem Phys 126:014101. 10.1063/1.2408420

10. Cagiada M, Thomasen FE, Ovchinnikov S, et al (2025) AF2χ: Predicting protein side-chain rotamer distributions with AlphaFold2. 2025.04.16.649219

11. Capdevila DA, Huerta F, Edmonds KA, et al (2018) Tuning site-specific dynamics to drive allosteric activation in a pneumococcal zinc uptake regulator. eLife 7:e37268. 10.7554/eLife.37268

12. Carbonell P, Sol A del (2009) Methyl side-chain dynamics prediction based on protein structure. Bioinformatics 25:2552–2558. 10.1093/bioinformatics/btp463

13. Curtis WA, Wenz J, Krüger CR, et al (2026) Ultrathin liquid cells for microsecond time-resolved cryo-EM. Nat Commun 17:1799. 10.1038/s41467-026-68515-z

14. Delaglio F, Grzesiek S, Vuister GW, et al (1995) NMRPipe: a multidimensional spectral processing system based on UNIX pipes. J Biomol NMR 6:277–293. 10.1007/BF00197809

15. Doig AJ, Sternberg MJ (1995) Side-chain conformational entropy in protein folding. Protein Sci 4:2247–2251. 10.1002/pro.5560041101

16. Dubey A, Stoyanov N, Viennet T, et al (2021) Local Deuteration Enables NMR Observation of Methyl Groups in Proteins from Eukaryotic and Cell-Free Expression Systems. Angewandte Chemie International Edition 60:13783–13787. 10.1002/anie.202016070

17. Eisenhaber F, Lijnzaad P, Argos P, et al (1995) The double cubic lattice method: Efficient approaches to numerical integration of surface area and volume and to dot surface contouring of molecular assemblies. Journal of Computational Chemistry 16:273–284. 10.1002/jcc.540160303

18. Frank J (2017) Time-resolved cryo-electron microscopy: Recent progress. Journal of Structural Biology 200:303–306. 10.1016/j.jsb.2017.06.005

19. Frederick KK, Marlow MS, Valentine KG, Wand AJ (2007) Conformational entropy in molecular recognition by proteins. Nature 448:325–329. 10.1038/nature05959

20. Gu Y, Li D-W, Brüschweiler R (2014) NMR Order Parameter Determination from Long Molecular Dynamics Trajectories for Objective Comparison with Experiment. J Chem Theory Comput 10:2599–2607. 10.1021/ct500181v

21. Halle B, Wennerström H (1981) Interpretation of magnetic resonance data from water nuclei in heterogeneous systems. J Chem Phys 75:1928–1943. 10.1063/1.442218

22. Han J, van Hoof A (2016) The RNA exosome channeling and direct access conformations have distinct in vivo functions. Cell Rep 16:3348–3358. 10.1016/j.celrep.2016.08.059

23. Henry ER, Szabo A (1985) Influence of vibrational motion on solid state line shapes and NMR relaxation. J Chem Phys 82:4753–4761. 10.1063/1.448692

24. Hoffmann F, Mulder FAA, Schäfer LV (2018a) Accurate Methyl Group Dynamics in Protein Simulations with AMBER Force Fields. J Phys Chem B 122:5038–5048. 10.1021/acs.jpcb.8b02769

25. Hoffmann F, Mulder FAA, Schäfer LV (2020) Predicting NMR relaxation of proteins from molecular dynamics simulations with accurate methyl rotation barriers. J Chem Phys 152:084102. 10.1063/1.5135379

26. Hoffmann F, Xue M, Schäfer LV, Mulder FAA (2018b) Narrowing the gap between experimental and computational determination of methyl group dynamics in proteins. Phys Chem Chem Phys 20:24577–24590. 10.1039/C8CP03915A

27. Hoover WG (1985) Canonical dynamics: Equilibrium phase-space distributions. Phys Rev A 31:1695–1697. 10.1103/PhysRevA.31.1695

28. Jorgensen WL, Maxwell DS, Tirado-Rives J (1996) Development and testing of the OPLS all-atom force field on conformational energetics and properties of organic liquids. J Am Chem Soc 118:11225–11236. 10.1021/ja9621760

29. Kasinath V, Sharp KA, Wand AJ (2013) Microscopic Insights into the NMR Relaxation-Based Protein Conformational Entropy Meter. J Am Chem Soc 135:15092–15100. 10.1021/ja405200u

30. Kay LE (2011) Solution NMR spectroscopy of supra-molecular systems, why bother? A methyl-TROSY view. J Magn Reson 210:159–170. 10.1016/j.jmr.2011.03.008

31. Kay LE, Prestegard JH (1987) Methyl group dynamics from relaxation of double quantum filtered NMR signals. Application to deoxycholate. J Am Chem Soc 109:3829–3835. 10.1021/ja00247a002

32. Kleckner IR, Foster MP (2011) An introduction to NMR-based approaches for measuring protein dynamics. Biochim Biophys Acta 1814:942–968. 10.1016/j.bbapap.2010.10.012

33. Kümmerer F, Orioli S, Harding-Larsen D, et al (2021) Fitting Side-Chain NMR Relaxation Data Using Molecular Simulations. J Chem Theory Comput 17:5262–5275. 10.1021/acs.jctc.0c01338

34. Liao X, Long D, Li D-W, et al (2012) Probing Side-Chain Dynamics in Proteins by the Measurement of Nine Deuterium Relaxation Rates Per Methyl Group. J Phys Chem B 116:606–620. 10.1021/jp209304c

35. Liebau J, Lazzaretti D, Fürtges T, et al (2025) 4D structural biology–quantitative dynamics in the eukaryotic RNA exosome complex. Nat Commun 16:7896. 10.1038/s41467-025-62982-6

36. Lindorff-Larsen K, Maragakis P, Piana S, Shaw DE (2016) Picosecond to Millisecond Structural Dynamics in Human Ubiquitin. J Phys Chem B 120:8313–8320. 10.1021/acs.jpcb.6b02024

37. Lipari G, Szabo A (1982a) Model-free approach to the interpretation of nuclear magnetic resonance relaxation in macromolecules. 1. Theory and range of validity. J Am Chem Soc 104:4546–4559. 10.1021/ja00381a009

38. Lipari G, Szabo A (1982b) Model-free approach to the interpretation of nuclear magnetic resonance relaxation in macromolecules. 2. Analysis of experimental results. J Am Chem Soc 104:4559–4570. 10.1021/ja00381a010

39. Lipari G, Szabo A, Levy RM (1982) Protein dynamics and NMR relaxation: comparison of simulations with experiment. Nature 300:197–198. 10.1038/300197a0

40. Liu Q, Greimann JC, Lima CD (2006) Reconstitution, activities, and structure of the eukaryotic RNA exosome. Cell 127:1223–1237. 10.1016/j.cell.2006.10.037

41. Lorentzen E, Dziembowski A, Lindner D, et al (2007) RNA channelling by the archaeal exosome. EMBO Rep 8:470–476. 10.1038/sj.embor.7400945

42. Makino DL, Baumgärtner M, Conti E (2013) Crystal structure of an RNA-bound 11-subunit eukaryotic exosome complex. Nature 495:70–75. 10.1038/nature11870

43. Makino DL, Schuch B, Stegmann E, et al (2015) RNA degradation paths in a 12-subunit nuclear exosome complex. Nature 524:54–58. 10.1038/nature14865

44. Markwick PRL, Bouvignies G, Blackledge M (2007) Exploring Multiple Timescale Motions in Protein GB3 Using Accelerated Molecular Dynamics and NMR Spectroscopy. J Am Chem Soc 129:4724–4730. 10.1021/ja0687668

45. Marlow MS, Dogan J, Frederick KK, et al (2010) The role of conformational entropy in molecular recognition by calmodulin. Nat Chem Biol 6:352–358. 10.1038/nchembio.347

46. Ming D, Brüschweiler R (2004) Prediction of methyl-side Chain Dynamics in Proteins. J Biomol NMR 29:363–368. 10.1023/B:JNMR.0000032612.70767.35

47. Mittermaier A, Davidson AR, Kay LE (2003) Correlation between 2H NMR Side-Chain Order Parameters and Sequence Conservation in Globular Proteins. J Am Chem Soc 125:9004–9005. 10.1021/ja034856q

48. Nosé S (1984) A unified formulation of the constant temperature molecular dynamics methods. J Chem Phys 81:511–519. 10.1063/1.447334

49. O’Brien ES, Wand AJ, Sharp KA (2016) On the ability of molecular dynamics force fields to recapitulate NMR derived protein side chain order parameters. Protein Science 25:1156–1160. 10.1002/pro.2922

50. Overbeck JH, Stelzig D, Fuchs A-L, et al (2022) Observation of conformational changes that underlie the catalytic cycle of Xrn2. Nat Chem Biol 18:1152–1160. 10.1038/s41589-022-01111-6

51. Parrinello M, Rahman A (1981) Polymorphic transitions in single crystals: A new molecular dynamics method. J Appl Phys 52:7182–7190. 10.1063/1.328693

52. Pintilie G, Shao C, Wang Z, et al (2025) Q-score as a reliability measure for protein, nucleic acid and small-molecule atomic coordinate models derived from 3DEM maps. Acta Cryst D 81:410–422. 10.1107/S2059798325005923

53. Pintilie G, Zhang K, Su Z, et al (2020) Measurement of atom resolvability in cryo-EM maps with Q-scores. Nat Methods 17:328–334. 10.1038/s41592-020-0731-1

54. Sapienza PJ, Lee AL (2010) Using NMR to study fast dynamics in proteins: methods and applications. Current Opinion in Pharmacology 10:723–730. 10.1016/j.coph.2010.09.006

55. Scherlo M, Wippermann E, Fürtges T, et al (2026) kontakteUR: transforming coordinates to chemical intuition to focus on essential interactions in biomolecular systems. 2026.06.19.732925

56. Schuller JM, Falk S, Fromm L, et al (2018) Structure of the nuclear exosome captured on a maturing preribosome. Science 360:219–222. 10.1126/science.aar5428

57. Schütz S, Sprangers R (2020) Methyl TROSY spectroscopy: A versatile NMR approach to study challenging biological systems. Prog Nucl Mag Res Sp 116:56–84

58. Sharp KA, Kasinath V, Wand AJ (2014) Banding of NMR-derived Methyl Order Parameters: Implications for Protein Dynamics. Proteins 82:2106–2117. 10.1002/prot.24566

59. Sheppard D, Sprangers R, Tugarinov V (2010) Experimental approaches for NMR studies of side-chain dynamics in high-molecular-weight proteins. Progress in Nuclear Magnetic Resonance Spectroscopy 56:1–45. 10.1016/j.pnmrs.2009.07.004

60. Sprangers R, Kay LE (2007) Quantitative dynamics and binding studies of the 20S proteasome by NMR. Nature 445:618–622. 10.1038/nature05512

61. Sun H, Kay LE, Tugarinov V (2011) An Optimized Relaxation-Based Coherence Transfer NMR Experiment for the Measurement of Side-Chain Order in Methyl-Protonated, Highly Deuterated Proteins. J Phys Chem B 115:14878–14884. 10.1021/jp209049k

62. Tugarinov V, Hwang PM, Ollerenshaw JE, Kay LE (2003) Cross-correlated relaxation enhanced ^1^H^13^C NMR spectroscopy of methyl groups in very high molecular weight proteins and protein complexes. J Am Chem Soc 125:10420–10428. 10.1021/ja030153x

63. Tugarinov V, Kay LE (2006) Relaxation Rates of Degenerate 1H Transitions in Methyl Groups of Proteins as Reporters of Side-Chain Dynamics. J Am Chem Soc 128:7299–7308. 10.1021/ja060817d

64. Tugarinov V, Ollerenshaw JE, Kay LE (2005) Probing Side-Chain Dynamics in High Molecular Weight Proteins by Deuterium NMR Spin Relaxation: An Application to an 82-kDa Enzyme. J Am Chem Soc 127:8214–8225. 10.1021/ja0508830

65. Tugarinov V, Sprangers R, Kay LE (2007) Probing Side-Chain Dynamics in the Proteasome by Relaxation Violated Coherence Transfer NMR Spectroscopy. J Am Chem Soc 129:1743–1750. 10.1021/ja067827z

66. Voss JM, Harder OF, Olshin PK, et al (2021) Rapid melting and revitrification as an approach to microsecond time-resolved cryo-electron microscopy. Chemical Physics Letters 778:138812. 10.1016/j.cplett.2021.138812

67. Wang Y, V. S. M, Kim J, et al (2019) Globally correlated conformational entropy underlies positive and negative cooperativity in a kinase’s enzymatic cycle. Nat Commun 10:799. 10.1038/s41467-019-08655-7

68. Wasmuth EV, Lima CD (2012) Exo- and endoribonucleolytic activities of yeast cytoplasmic and nuclear RNA exosomes are dependent on the noncatalytic core and central channel. Mol Cell 48:133–144. 10.1016/j.molcel.2012.07.012

69. Xu Y, Huang J (2021) Validating the CHARMM36m protein force field with LJ-PME reveals altered hydrogen bonding dynamics under elevated pressures. Commun Chem 4:99. 10.1038/s42004-021-00537-8

70. Yang D, Kay LE (1996) Contributions to Conformational Entropy Arising from Bond Vector Fluctuations Measured from NMR-Derived Order Parameters: Application to Protein Folding. Journal of Molecular Biology 263:369–382. 10.1006/jmbi.1996.0581

71. Zinder JC, Lima CD (2017) Targeting RNA for processing or destruction by the eukaryotic RNA exosome and its cofactors. Genes Dev 31:88–100. 10.1101/gad.294769.116

72. Zumpfe K, Berbon M, Habenstein B, et al (2024) Analytical Framework to Understand the Origins of Methyl Side-Chain Dynamics in Protein Assemblies. J Am Chem Soc 146:8164–8178. 10.1021/jacs.3c12620

