## Supplementary Material for "Nanosecond methyl dynamics in the eukaryotic RNA exosome core"

### Table of contents

Supplementary Tables S1-S6 Page S2 – S11

Supplementary Figures S1-S11 Page S12 – S35

**Table S1: *C. thermophilum* proteins employed in this study with Uniprot and internal database reference.**

<table>
<tr><th>Protein</th><th>Accession code</th><th>Internal database reference</th></tr>
<tr><td>Rrp41</td><td>G0SC21</td><td>#1245</td></tr>
<tr><td>Rrp45</td><td>G0S755</td><td>#1215</td></tr>
<tr><td>Rrp43</td><td>G0S1P1</td><td rowspan="2">#1353</td></tr>
<tr><td>Rrp46</td><td>G0SCD1</td></tr>
<tr><td>Rrp42</td><td>G0RZG4</td><td>#1214</td></tr>
<tr><td>Mtr3</td><td>P0CT46</td><td>#1246</td></tr>
<tr><td>Rrp40</td><td>G0RZX8</td><td>#1213</td></tr>
<tr><td>Rrp4</td><td>G0S9A0</td><td>#1233</td></tr>
<tr><td>Csl4</td><td>G0SE33</td><td>#1247</td></tr>
</table>

**Table S2: Sequence of RNA employed in this study.**

| RNA | Sequence | Internal database reference |
|---|---|---|
| 30mer | GGAGGAGAGGAAGGUGGAAGAAAGAAGAGG | #73 |

**Table S3: (A) Assignment mutants of Rrp40 with internal database reference and primers used for construct design. (B) Ile-δ1 methyl group assignment for Rrp40 monomer and when reconstituted into Exo9.**

**(A)**

| Construct | Internal database reference | forward primer | reverse primer |
|---|---|---|---|
| Rrp40-I15V | 2791 | CTGCCGGGCGAAACCGTTGATCCGAGTCTGG | CCAGACTCGGATCAACGGTTTCGCCCGGCAG |
| Rrp40-I42V | 2798 | GTTCCGCCGAGCGATGTTATCCCGACCGTGG | CCACGGTCGGGATAACATCGCTCGGCGGAAC |
| Rrp40-I43V | 2799 | GTTCCGCCGAGCGATATTGTCCCGACCGTGGC | GCCACGGTCGGGACAATATCGCTCGGCGGAAC |
| Rrp40-I51V | 2792 | GTGGCCGGCCAGCTGGTTACGAACCTGAAC | GTTCAGGTTCGTAACCAGCTGGCCGGCCAC |
| Rrp40-I92V | 2796 | CTCTTATCTGTGTCTGGTCACCCCGCATACGC | GCGTATGCGGGGTGACCAGACACAGATAAGAG |
| Rrp40-I154V | 2797 | GATGGTCTGGGTCCGGTTACCGGTCCGGGTTG | CAACCCGGACCGGTAACCGGACCCAGACCATC |
| Rrp40-I195V | 2793 | GCGAAGATCCGAGTGTTGGTGAAGCCGGTGC | GCACCGGCTTCACCAACACTCGGATCTTCGC |
| Rrp40-I225V | 2794 | GATGTGAAAACCGTTGTTATCGTCGGTCGTGCACTGC | GCAGTGCACGACCGACGATAACAACGGTTTTCACATC |
| Rrp40-I226V | 2800 | GTGAAAACCGTTATTGTCGTCGGTCGTGCACTGC | GCAGTGCACGACCGACGACAATAACGGTTTTCAC |
| Rrp40-I241V | 2795 | GACCGCGGCAACCTGACGGTTGAAGGTCAACGC | GCGTTGACCTTCAACCGTCAGGTTGCCGCGGTC |

**(B)**

| Resonance | | monomer | | Exo9 | |
|---|---|---|---|---|---|
| | | $^{1}H$ (ppm) | $^{13}C$ (ppm) | $^{1}H$ (ppm) | $^{13}C$ (ppm) |
| Rrp40-I15[1] | | ~0.906 | ~10.21 | 0.810 | 9.34 |
| Rrp40-I42[1] | | ~0.910 | ~10.16 | 0.899 | 10.76 |
| Rrp40-I43 | | 0.922 | 9.88 | 0.871 | 11.03 |
| Rrp40-I51[1] | | ~0.912 | ~10.12 | 0.547 | 7.74 |
| Rrp40-I92 | | 0.979 | 10.45 | 1.030 | 10.47 |
| Rrp40-I154 | A | 0.248 | 8.06 | 0.130 | 7.79 |
| | B | 0.039 | 8.29 | 0.029 | 8.00 |
| Rrp40-I195 | | 0.928 | 9.44 | 0.923 | 9.27 |
| Rrp40-I225[1] | | ~0.905 | ~10.48 | 0.859 | 11.76 |
| Rrp40-I226 | | 0.790 | 10.51 | 0.802 | 10.84 |
| Rrp40-I241 | | 0.840 | 7.89 | 0.688 | 6.71 |

[1] Due to overlap the assignment is approximate in the monomer.

**Table S4: Protein constructs and NMR experiments conducted on them in this study.**

| Construct | Type(s) of experiment |
|---|---|
| I-labeled monomeric Rrp40 | Methyl TROSY HMQC |
| I-labeled monomeric Rrp40-I15V | Methyl TROSY HMQC |
| I-labeled monomeric Rrp40-I42V | Methyl TROSY HMQC |
| I-labeled monomeric Rrp40-I43V | Methyl TROSY HMQC |
| I-labeled monomeric Rrp40-I51V | Methyl TROSY HMQC |
| I-labeled monomeric Rrp40-I92V | Methyl TROSY HMQC |
| I-labeled monomeric Rrp40-I154V | Methyl TROSY HMQC |
| I-labeled monomeric Rrp40-I195V | Methyl TROSY HMQC |
| I-labeled monomeric Rrp40-I225V | Methyl TROSY HMQC |
| I-labeled monomeric Rrp40-I226V | Methyl TROSY HMQC |
| I-labeled monomeric Rrp40-I241V | Methyl TROSY HMQC |
| Exo9, I-labeled Rrp40 | Methyl TROSY HMQC, TQ forbidden experiment |
| Exo9, I-labeled Rrp40-I15V | Methyl TROSY HMQC |
| Exo9, I-labeled Rrp40-I42V | Methyl TROSY HMQC |
| Exo9, I-labeled Rrp40-I43V | Methyl TROSY HMQC |
| Exo9, I-labeled Rrp40-I51V | Methyl TROSY HMQC |
| Exo9, I-labeled Rrp40-I92V | Methyl TROSY HMQC |
| Exo9, I-labeled Rrp40-I154V | Methyl TROSY HMQC |
| Exo9, I-labeled Rrp40-I195V | Methyl TROSY HMQC |
| Exo9, I-labeled Rrp40-I225V | Methyl TROSY HMQC |
| Exo9, I-labeled Rrp40-I226V | Methyl TROSY HMQC |
| Exo9, I-labeled Rrp40-I241V | Methyl TROSY HMQC |
| Exo9, I-labeled Csl4 | Methyl TROSY HMQC, TQ forbidden experiment |
| Exo9, I-labeled Rrp41 | Methyl TROSY HMQC, TQ forbidden experiment |

| | |
|---|---|
| Exo9, I-labeled Rrp45 | Methyl TROSY HMQC, TQ forbidden experiment |
| Exo9 + 30mer RNA, I-labeled Csl4 | Methyl TROSY HMQC, TQ forbidden experiment |
| Exo9 + 30mer RNA, I-labeled Rrp40 | Methyl TROSY HMQC, TQ forbidden experiment |
| Exo9 + 30mer RNA, I-labeled Rrp41 | Methyl TROSY HMQC, TQ forbidden experiment |
| Exo9 + 30mer RNA, I-labeled Rrp45 | Methyl TROSY HMQC, TQ forbidden experiment |

**Table S5: Experimental methyl order parameters $S_{axis}^2$ and effective proton distances $r_{eff}$ for Csl4, Rrp40, Rrp41 and Rrp45 in Exo9 in the absence and presence of RNA and methyl order parameters $S_{axis}^2{}_{MD}$ calculated from the MD simulation.** Uncertainties are given ±1 SD. n.a.: resonance experimentally not observed.

| | no RNA | | with RNA | | no RNA | | with RNA | | no RNA | |
|---|---|---|---|---|---|---|---|---|---|---|
| Resonance | $S_{axis}^2$ | SD $S_{axis}^2$ | $S_{axis}^2$ | SD $S_{axis}^2$ | $r_{eff}$ (Å) | SD $r_{eff}$ (Å) | $r_{eff}$ (Å) | SD $r_{eff}$ (Å) | $S_{axis}^2{}_{MD}$ | SD $S_{axis}^2{}_{MD}$ |
| Csl4-I18 | 0.35 | 0.03 | 0.32 | 0.07 | 2.5 | 0.1 | 2.5 | 0.3 | 0.54 | 0.09 |
| Csl4-I60[1] | 0.035 | 0.002 | 0.039 | 0.002 | 3.3 | 0.2 | 3.2 | 0.3 | 0.21 | 0.16 |
| Csl4-I72 | 0.67 | 0.21 | 1.14 | 0.45 | 2.3 | 0.4 | 2.1 | 0.3 | 0.49 | 0.23 |
| Csl4-I95[1] | 0.035 | 0.002 | 0.039 | 0.002 | 3.3 | 0.2 | 3.2 | 0.3 | 0.58 | 0.24 |
| Csl4-I102 | 0.52 | 0.05 | 0.57 | 0.11 | 3.1 | 0.3 | 2.6 | 0.4 | 0.24 | 0.15 |
| Csl4-I108 | 0.34 | 0.04 | 0.52 | 0.16 | 2.5 | 0.1 | 2.3 | 0.1 | 0.76 | 0.08 |
| Csl4-I110 | 0.46 | 0.49 | 0.83 | 0.53 | 2.8 | 0.5 | 2.0 | 0.4 | 0.76 | 0.12 |
| Csl4-I119 | 0.96 | 0.41 | 1.00 | 0.49 | 2.2 | 0.3 | 2.3 | 0.5 | 0.73 | 0.1 |
| Csl4-I135 | 0.94 | 0.35 | 0.63 | 0.52 | 2.0 | 0.2 | 2.4 | 0.5 | 0.82 | 0.02 |
| Csl4-I140 | 1.26 | 0.48 | 0.84 | 0.40 | 2.0 | 0.3 | 2.5 | 0.6 | 0.76 | 0.09 |
| Csl4-I159 | 0.30 | 0.52 | 0.91 | 0.47 | 927.9 | 0.7 | 2.0 | 0.4 | 0.48 | 0.16 |
| Csl4-I165 | 0.66 | 0.37 | 0.90 | 0.56 | 2.5 | 0.5 | 2.3 | 0.6 | 0.82 | 0.05 |
| Csl4-I185 | 0.71 | 0.39 | 0.75 | 0.29 | 2.0 | 0.3 | 2.0 | 0.3 | 0.47 | 0.22 |
| Csl4-I207 | 0.28 | 0.01 | 0.31 | 0.04 | 3.2 | 0.2 | 3.0 | 0.4 | 0.31 | 0.18 |
| Rrp40-I15 | 0.37 | 0.05 | 0.29 | 0.03 | 2.7 | 0.2 | 2.8 | 0.2 | 0.39 | 0.22 |
| Rrp40-I42 | 0.40 | 0.05 | 0.36 | 0.05 | 2.7 | 0.3 | 2.7 | 0.3 | 0.43 | 0.21 |
| Rrp40-I43 | 0.66 | 0.16 | 0.47 | 0.08 | 2.3 | 0.2 | 2.6 | 0.3 | 0.44 | 0.26 |
| Rrp40-I51 | 0.49 | 0.10 | 0.42 | 0.08 | 2.3 | 0.2 | 2.4 | 0.3 | 0.51 | 0.15 |
| Rrp40-I92 | 0.91 | 0.34 | 0.58 | 0.27 | 2.2 | 0.4 | 2.5 | 0.5 | 0.69 | 0.09 |
| Rrp40-I154 | 0.24 | 0.02 | 0.19 | 0.02 | 3.7 | 0.5 | 4.3 | 0.5 | 0.22 | 0.13 |

| | | | | | | | | | | |
|---|---|---|---|---|---|---|---|---|---|---|
| Rrp40-I195 | 0.22 | 0.02 | 0.18 | 0.01 | 3.3 | 0.4 | 3.2 | 0.3 | 0.43 | 0.18 |
| Rrp40-I226 | 0.75 | 0.11 | 0.61 | 0.11 | 2.4 | 0.2 | 2.5 | 0.3 | 0.56 | 0.27 |
| Rrp40-I241 | 0.66 | 0.16 | 0.48 | 0.09 | 2.7 | 0.4 | 2.9 | 0.5 | 0.26 | 0.11 |
| Rrp41-I30 | 0.84 | 0.37 | 0.66 | 0.38 | 2.3 | 0.4 | 2.5 | 0.5 | 0.73 | 0.1 |
| Rrp41-I96 | 0.59 | 0.34 | 0.63 | 0.42 | 2.6 | 0.5 | 2.3 | 0.5 | 0.85 | 0.02 |
| Rrp41-I98 | 0.51 | 0.12 | 0.42 | 0.20 | 2.2 | 0.2 | 2.3 | 0.4 | 0.56 | 0.24 |
| Rrp41-I116 | 0.33 | 0.07 | 0.26 | 0.15 | 2.4 | 0.2 | 3.4 | 0.6 | 0.53 | 0.26 |
| Rrp41-I117[2] | 0.46 | 0.08 | 0.41 | 0.09 | 2.5 | 0.2 | 2.3 | 0.1 | 0.56 | 0.24 |
| Rrp41-I141 | 1.21 | 0.50 | 0.66 | 0.45 | 1.9 | 0.5 | 2.3 | 0.5 | 0.63 | 0.09 |
| Rrp41-I143 | 0.74 | 0.25 | 0.52 | 0.21 | 2.0 | 0.3 | 2.2 | 0.4 | 0.35 | 0.16 |
| Rrp41-I159 | 0.72 | 0.34 | 1.10 | 0.49 | 2.4 | 0.5 | 2.0 | 0.4 | 0.84 | 0.03 |
| Rrp41-I171 | 1.44 | 0.49 | 0.70 | 0.38 | 1.8 | 0.4 | 2.1 | 0.4 | 0.66 | 0.17 |
| Rrp41-I251[2] | 0.46 | 0.08 | 0.20 | 0.19 | 2.5 | 0.2 | 2.8 | 0.6 | 0.43 | 0.2 |
| Rrp41-I254 | 1.01 | 0.35 | 1.74 | 0.53 | 2.2 | 0.4 | 1.9 | 0.3 | 0.39 | 0.14 |
| Rrp41-I267 | 0.27 | 0.20 | 0.43 | 0.18 | 2.7 | 0.5 | 2.2 | 0.3 | 0.3 | 0.19 |
| Rrp45-I76 | 0.61 | 0.13 | 0.51 | 0.55 | 2.4 | 0.3 | 812.6 | 0.5 | 0.47 | 0.19 |
| Rrp45-I79 | 0.76 | 0.25 | 0.47 | 0.22 | 2.3 | 0.3 | 2.6 | 0.5 | 0.35 | 0.25 |
| Rrp45-I111 | 0.51 | 0.20 | 0.35 | 0.33 | 2.2 | 0.4 | 2.6 | 0.5 | 0.41 | 0.19 |
| Rrp45-I133 | 0.61 | 0.22 | 0.64 | 0.19 | 2.6 | 0.5 | 2.3 | 0.3 | 0.37 | 0.13 |
| Rrp45-I152 | 0.83 | 0.42 | 0.59 | 0.30 | 2.3 | 0.4 | 2.3 | 0.4 | 0.82 | 0.06 |
| Rrp45-I175 | 0.49 | 0.17 | 0.37 | 0.09 | 2.3 | 0.3 | 2.3 | 0.2 | 0.57 | 0.16 |
| Rrp45-I206[3] | 0.25 | 0.02 | 0.19 | 0.01 | 3.4 | 0.4 | 2.9 | 0.2 | 0.53 | 0.2 |
| Rrp45-I225 | 0.53 | 0.11 | 0.43 | 0.12 | 2.2 | 0.1 | 2.3 | 0.3 | 0.51 | 0.15 |
| Rrp45-I255 | 0.75 | 0.19 | 0.64 | 0.11 | 2.4 | 0.3 | 2.4 | 0.2 | 0.51 | 0.13 |

[1] Resonances Csl4-I60 and Csl4-I95 overlap.

[2] Resonances Rrp41-I117 and Rrp41-I251 overlap in absence of RNA. In the presence of RNA the assignment is ambiguous.

[3] Resonance Rrp45-I206 overlaps with one or several unassigned resonances.

**Table S6: Spearman's rank correlation coefficient ρ, *p*-value between $S_{axis}^2$ and other parameters and the maximum expected ρ, given the measurement uncertainties and distribution of $S_{axis}^2$, $\rho_{ceil}$.** Uncertainties for ρ and $\rho_{ceil}$ are given ±1 SD and *p*-values were evaluated using a two-sided paired-sample *t*-test.

| Structural parameter | ρ | SD ρ | *p*-value | $\rho_{ceil}$ | SD $\rho_{ceil}$ |
|---|---|---|---|---|---|
| SASA | -0.47 | 0.10 | 0.00130 | 0.72 | 0.07 |
| RMSF | -0.44 | 0.11 | 0.0026 | 0.72 | 0.07 |
| $S_{axis}^2{}_{MD}$ | 0.38 | 0.13 | 0.011 | 0.72 | 0.07 |
| AF2$\chi_{single}$ (all subunits) | 0.46 | 0.10 | 0.003 | 0.72 | 0.07 |
| AF2$\chi_{complex}$ (all subunits) | 0.50 | 0.10 | 0.001 | 0.72 | 0.07 |
| AF2$\chi_{single}$ (Csl4) | 0.27 | 0.24 | 0.418 | 0.63 | 0.19 |
| AF2$\chi_{complex}$ (Csl4) | 0.32 | 0.21 | 0.341 | 0.63 | 0.19 |
| AF2$\chi_{single}$ (Rrp40) | 0.62 | 0.19 | 0.086 | 0.86 | 0.07 |
| AF2$\chi_{complex}$ (Rrp40) | 0.82 | 0.13 | 0.011 | 0.86 | 0.07 |
| AF2$\chi_{single}$ (Rrp41) | 0.32 | 0.18 | 0.317 | 0.75 | 0.11 |
| AF2$\chi_{complex}$ (Rrp41) | 0.40 | 0.21 | 0.198 | 0.75 | 0.11 |
| AF2$\chi_{single}$ (Rrp45) | 0.29 | 0.32 | 0.501 | 0.49 | 0.22 |
| AF2$\chi_{complex}$ (Rrp45) | 0.55 | 0.32 | 0.171 | 0.49 | 0.22 |
| X-ray resolution residue | -0.36 | 0.10 | 0.019 | 0.74 | 0.06 |
| X-ray resolution backbone | -0.36 | 0.10 | 0.018 | 0.74 | 0.06 |
| X-ray resolution side chain | -0.35 | 0.10 | 0.025 | 0.74 | 0.06 |
| cryo-EM resolution residue | -0.29 | 0.10 | 0.060 | 0.74 | 0.06 |
| cryo-EM resolution backbone | -0.30 | 0.10 | 0.056 | 0.74 | 0.06 |
| cryo-EM resolution side chain | -0.27 | 0.10 | 0.089 | 0.74 | 0.06 |
| cryo-EM Q-factor residue | 0.42 | 0.10 | 0.005 | 0.74 | 0.06 |
| cryo-EM Q-factor backbone | 0.19 | 0.09 | 0.222 | 0.74 | 0.06 |
| cryo-EM Q-factor side chain | 0.40 | 0.09 | 0.009 | 0.74 | 0.06 |

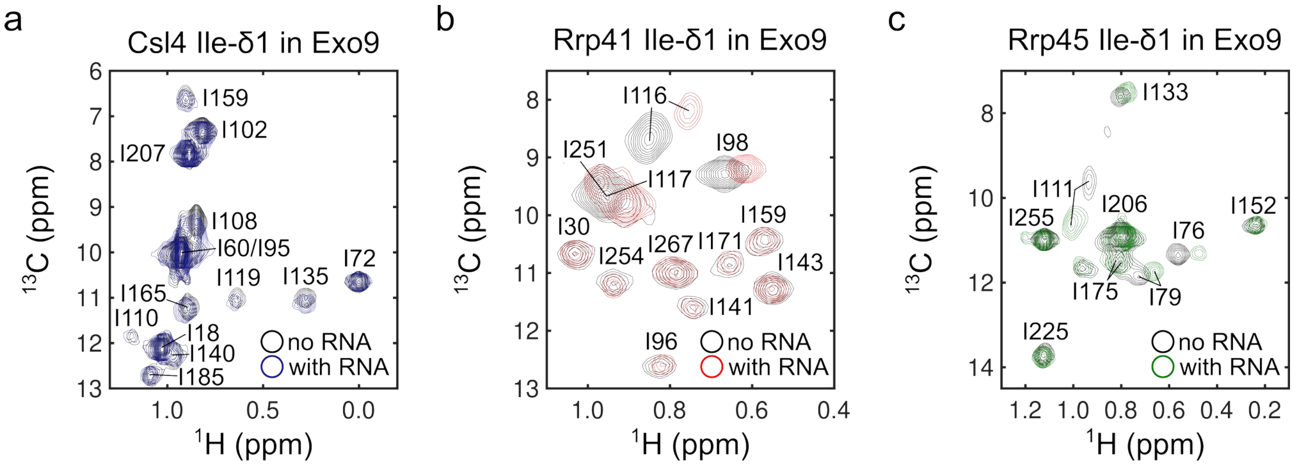


**Figure S1: HMQC of the Ile-δ1 region of (a) Csl4, (b) Rrp41, and (c) Rrp45 in the absence (black) and presence (colored) of RNA.**

a

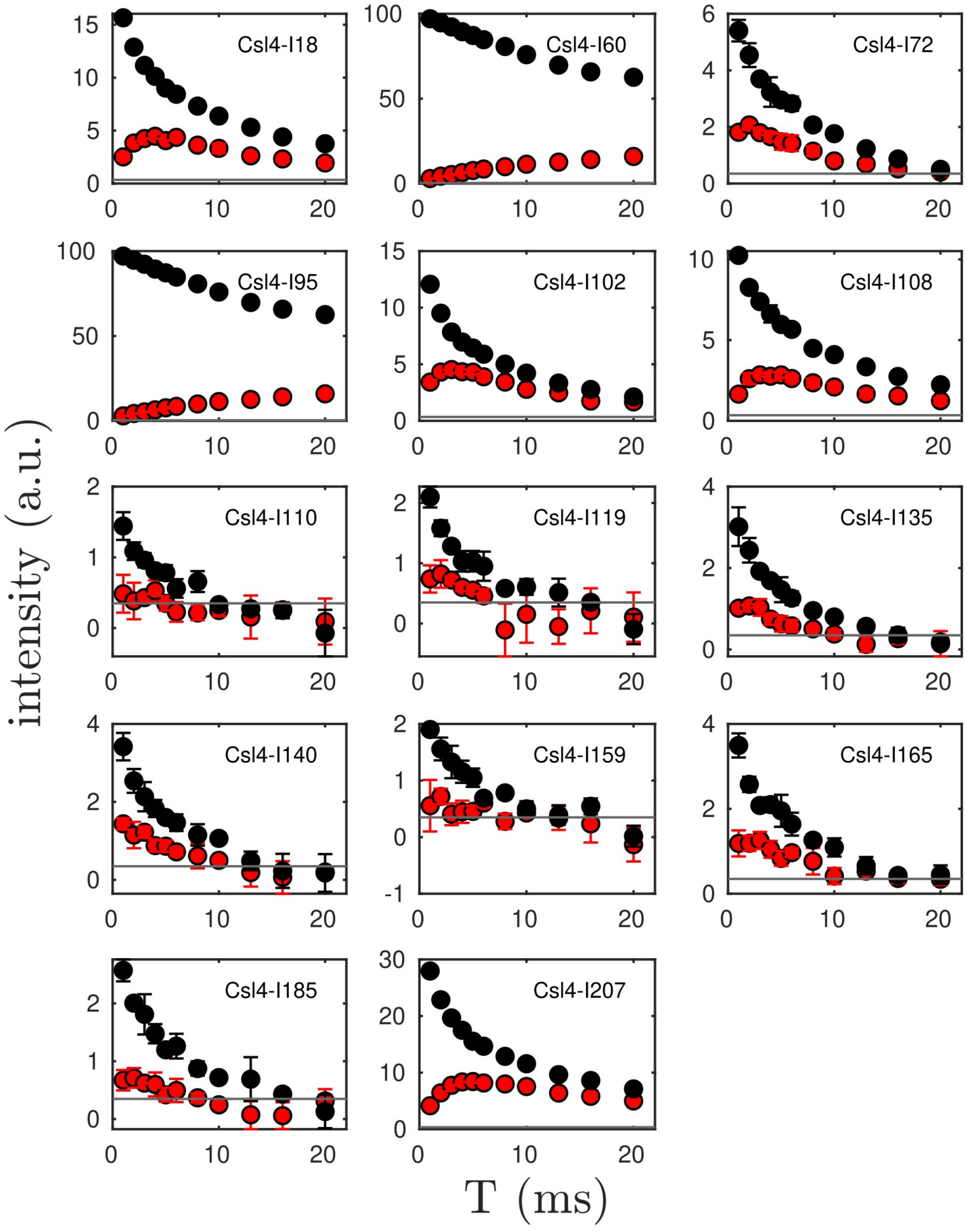
Csl4-I18
Csl4-I60
Csl4-I72
Csl4-I95
Csl4-I102
Csl4-I108
Csl4-I110
Csl4-I119
Csl4-I135
Csl4-I140
Csl4-I159
Csl4-I165
Csl4-I185
Csl4-I207
intensity (a.u.)
T (ms)

64

b

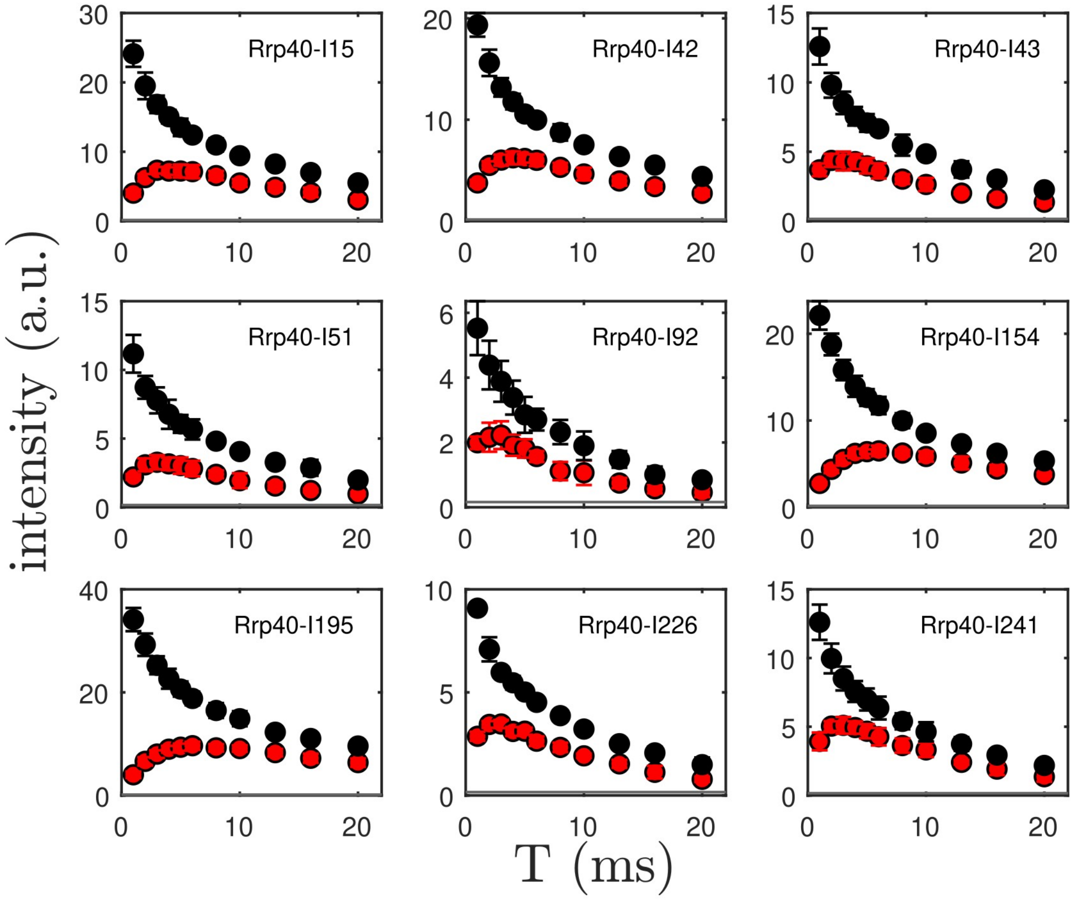
Rrp40-I15
Rrp40-I42
Rrp40-I43
Rrp40-I51
Rrp40-I92
Rrp40-I154
Rrp40-I195
Rrp40-I226
Rrp40-I241
intensity (a.u.)
T (ms)

C

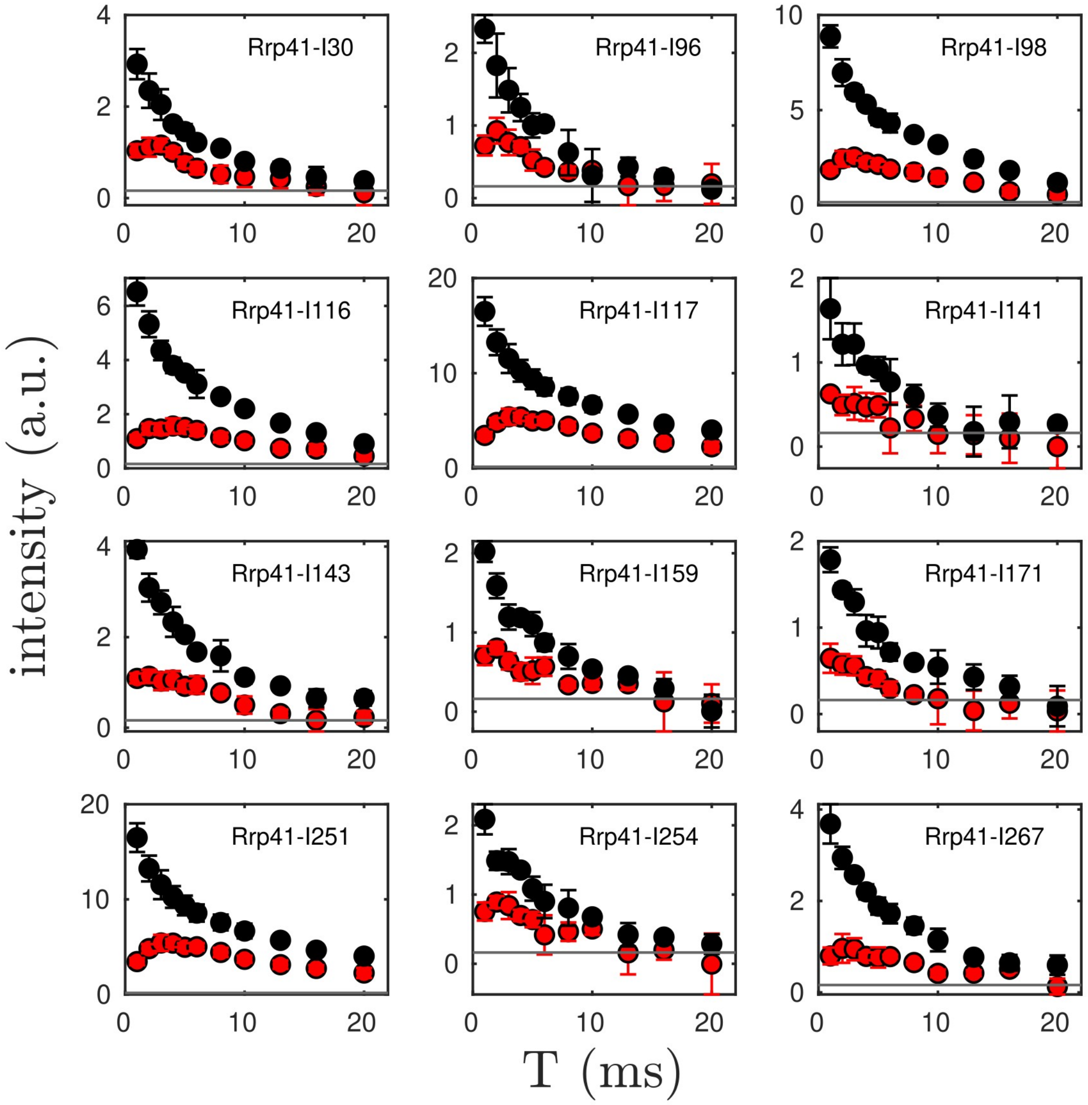
Rrp41-I30
Rrp41-I96
Rrp41-I98
Rrp41-I116
Rrp41-I117
Rrp41-I141
Rrp41-I143
Rrp41-I159
Rrp41-I171
Rrp41-I251
Rrp41-I254
Rrp41-I267
intensity (a.u.)
T (ms)

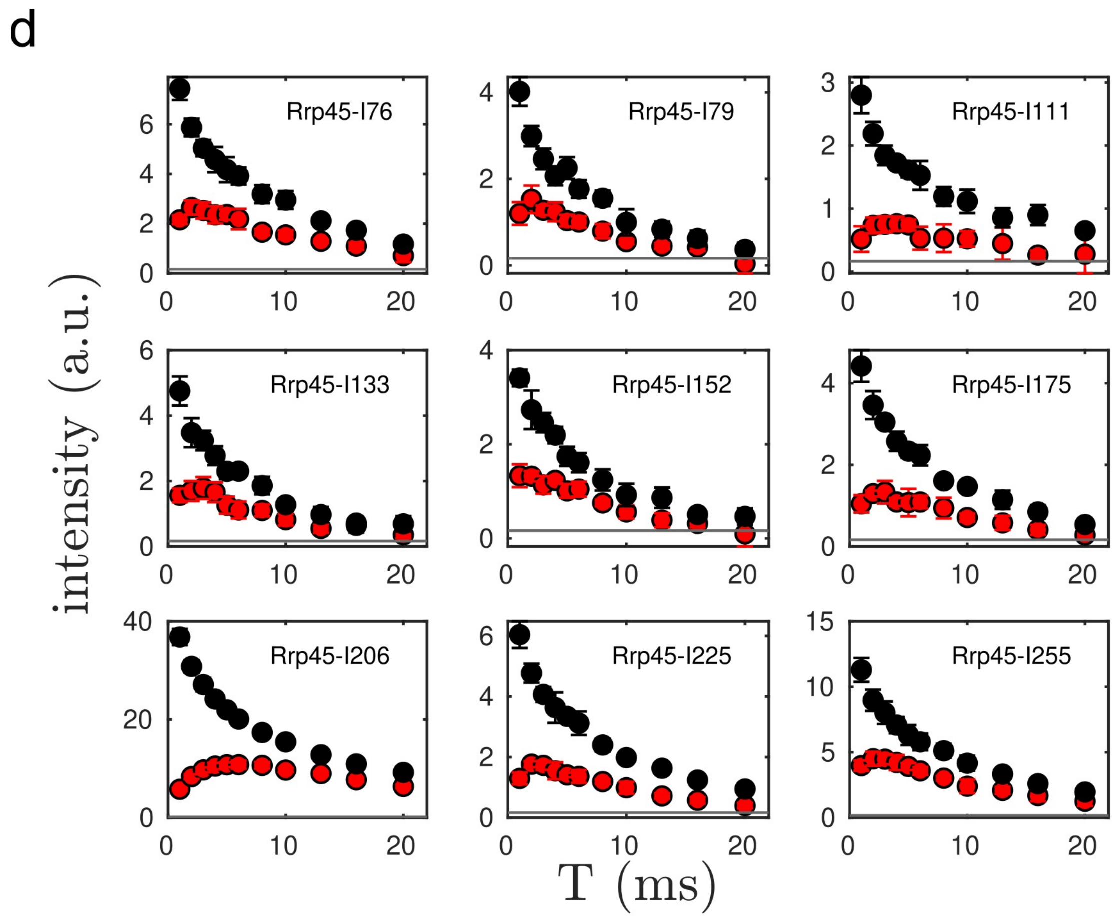


**Figure S2: SQ decay (black) and TQ build-up (red) for Ile-δ1 methyl groups in (a) Csl4, (b) Rrp40, (c) Rrp41 and (d) Rrp45 reconstituted into Exo9.** The horizontal grey line indicates the noise level. Data below the noise level was not considered in subsequent analyses. Note that the assignment for residues Csl4-I60/Csl4-I95, Rrp41-I117/Rrp41-I251 and Rrp45-I206 are ambiguous. Csl4-I60 and Csl4-I95 as well as Rrp41-I117 and Rrp41-I251 overlap and Rrp45-I206 overlaps with one or more unassigned resonance.

a

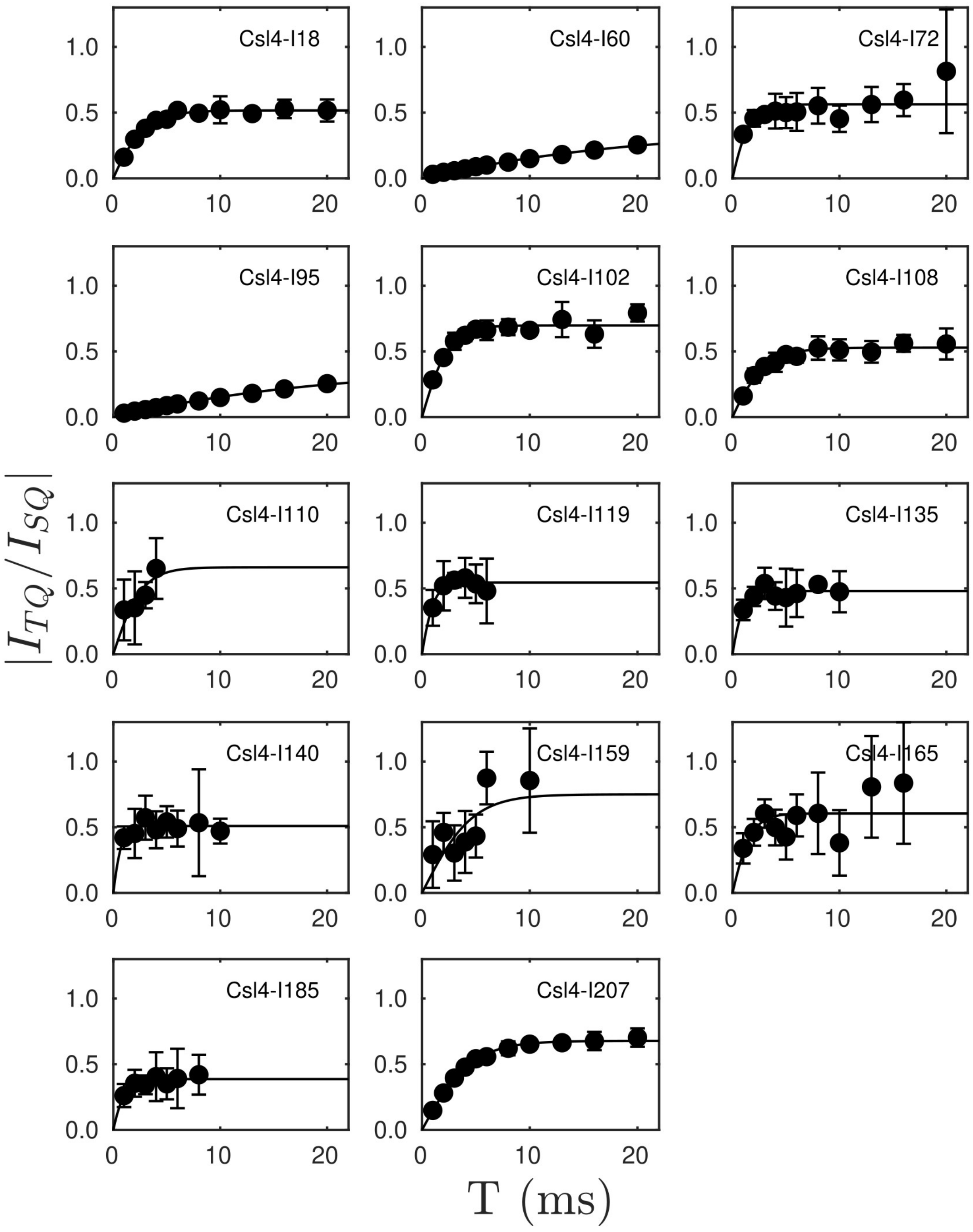
Csl4-I18
Csl4-I60
Csl4-I72
Csl4-I95
Csl4-I102
Csl4-I108
Csl4-I110
Csl4-I119
Csl4-I135
Csl4-I140
Csl4-I159
Csl4-I165
Csl4-I185
Csl4-I207
$|I_{TQ}/I_{SQ}|$
T (ms)

b

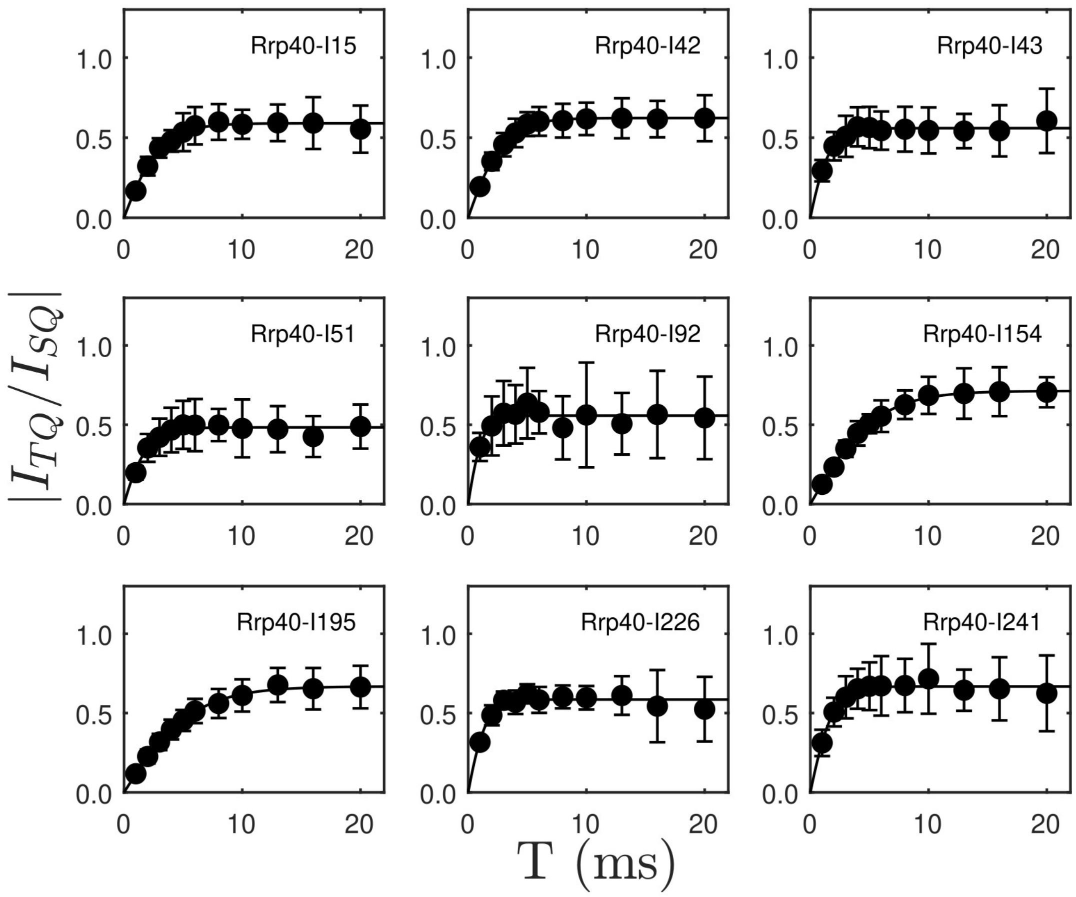
Rrp40-I15
Rrp40-I42
Rrp40-I43
Rrp40-I51
Rrp40-I92
Rrp40-I154
Rrp40-I195
Rrp40-I226
Rrp40-I241
$|I_{TQ}/I_{SQ}|$
T (ms)

C

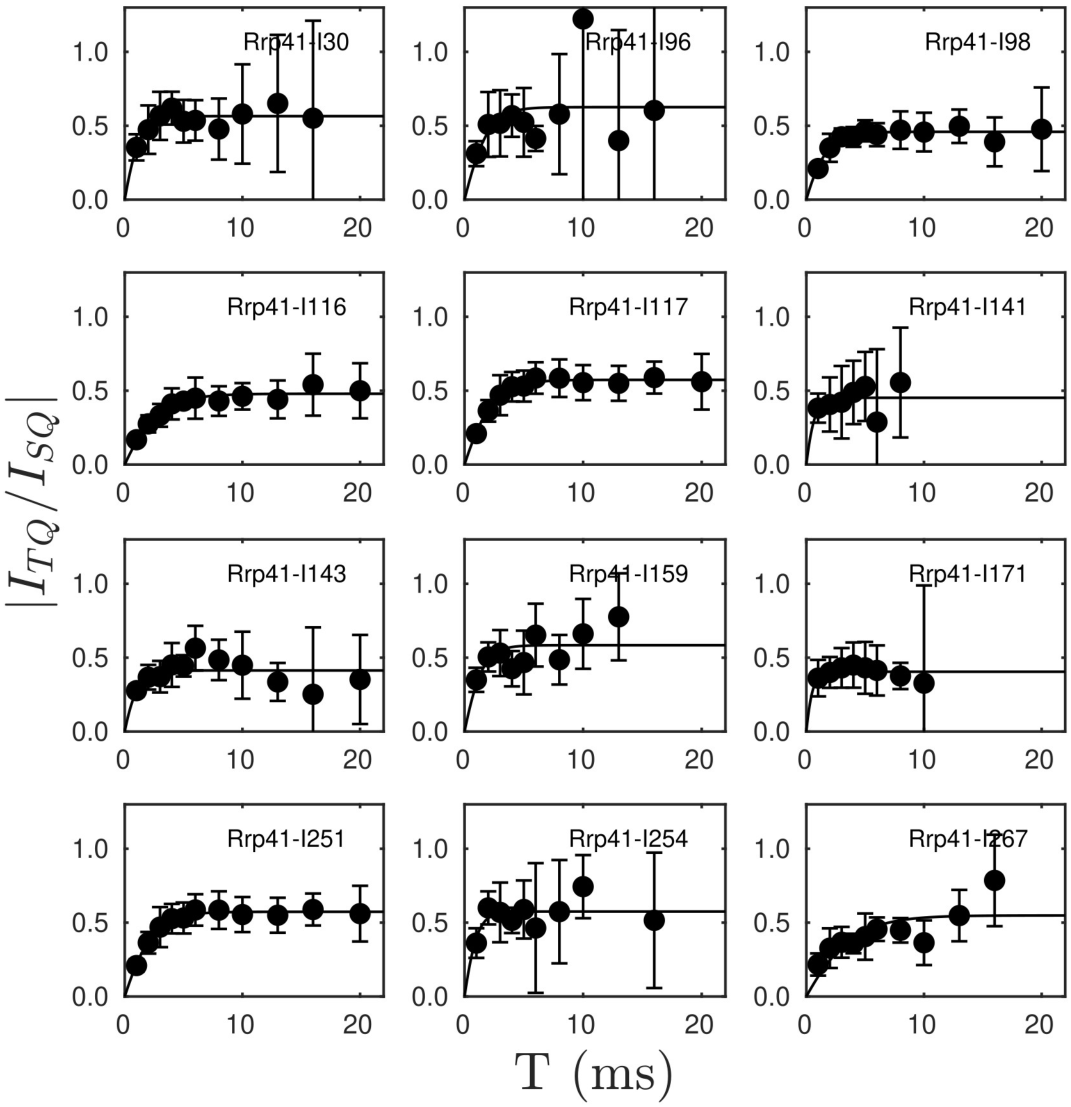

Rrp41-I30
Rrp41-I96
Rrp41-I98
Rrp41-I116
Rrp41-I117
Rrp41-I141
Rrp41-I143
Rrp41-I159
Rrp41-I171
Rrp41-I251
Rrp41-I254
Rrp41-I267
$|I_{TQ}/I_{SQ}|$
T (ms)

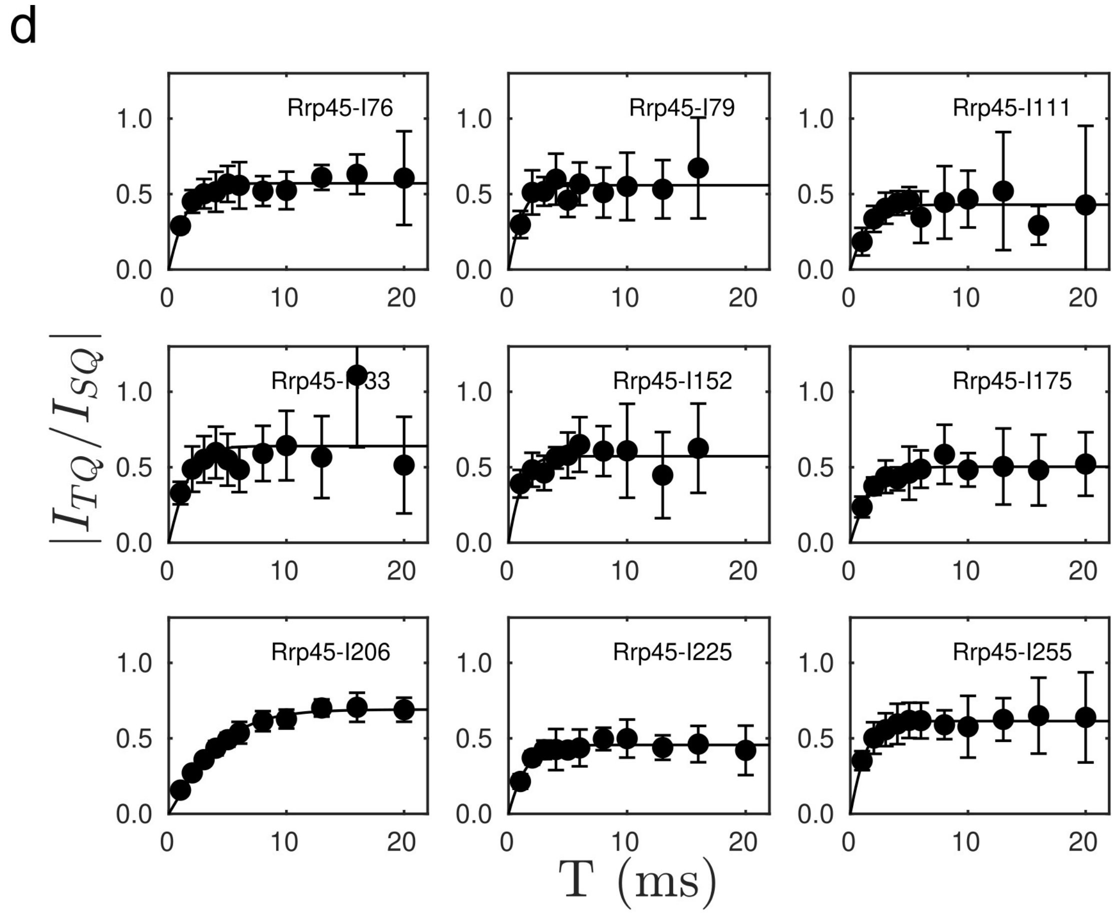


**Figure S3: Order parameter build-up curves for Ile-δ1 methyl groups in (a) Csl4, (b) Rrp40, (c) Rrp41 and (d) Rrp45 reconstituted into Exo9.** Note that the assignment for residues Csl4-I60/Csl4-I95, Rrp41-I117/Rrp41-I251 and Rrp45-I206 are ambiguous. Csl4-I60 and Csl4-I95 as well as Rrp41-I117 and Rrp41-I251 overlap and Rrp45-I206 overlaps with one or more unassigned resonance.

91 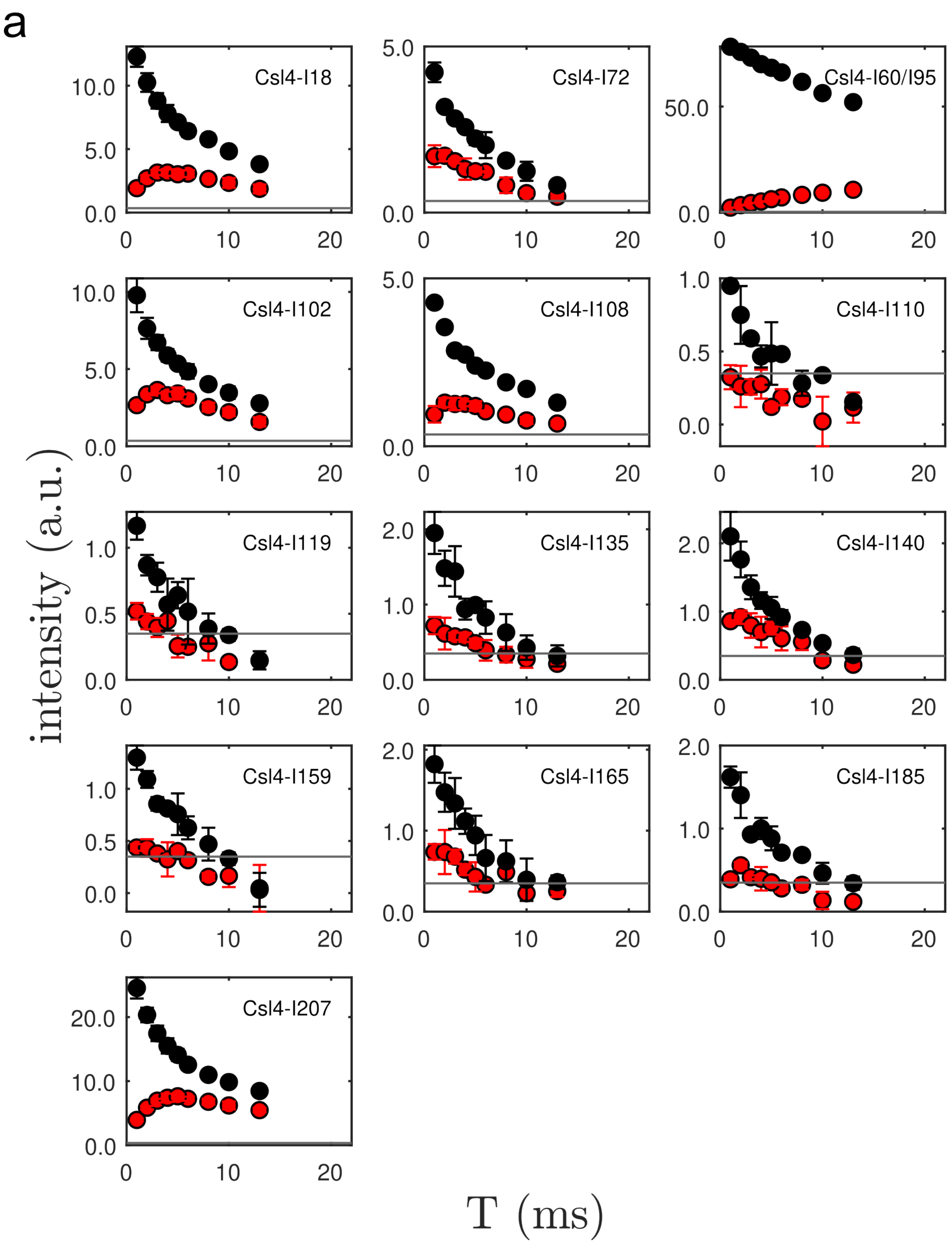

a
Csl4-I18
Csl4-I72
Csl4-I60/I95
Csl4-I102
Csl4-I108
Csl4-I110
Csl4-I119
Csl4-I135
Csl4-I140
Csl4-I159
Csl4-I165
Csl4-I185
Csl4-I207
intensity (a.u.)
T (ms)

b

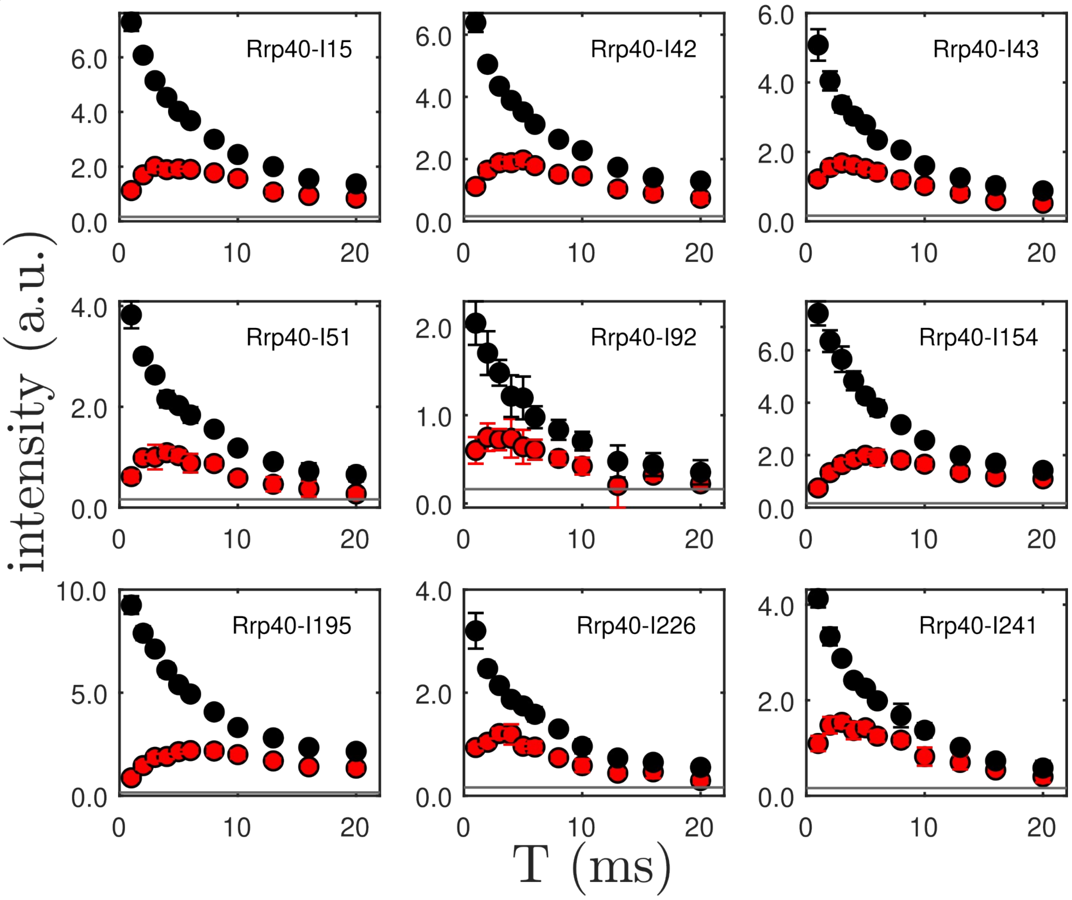

Rrp40-I15
Rrp40-I42
Rrp40-I43
Rrp40-I51
Rrp40-I92
Rrp40-I154
Rrp40-I195
Rrp40-I226
Rrp40-I241
intensity (a.u.)
T (ms)

c

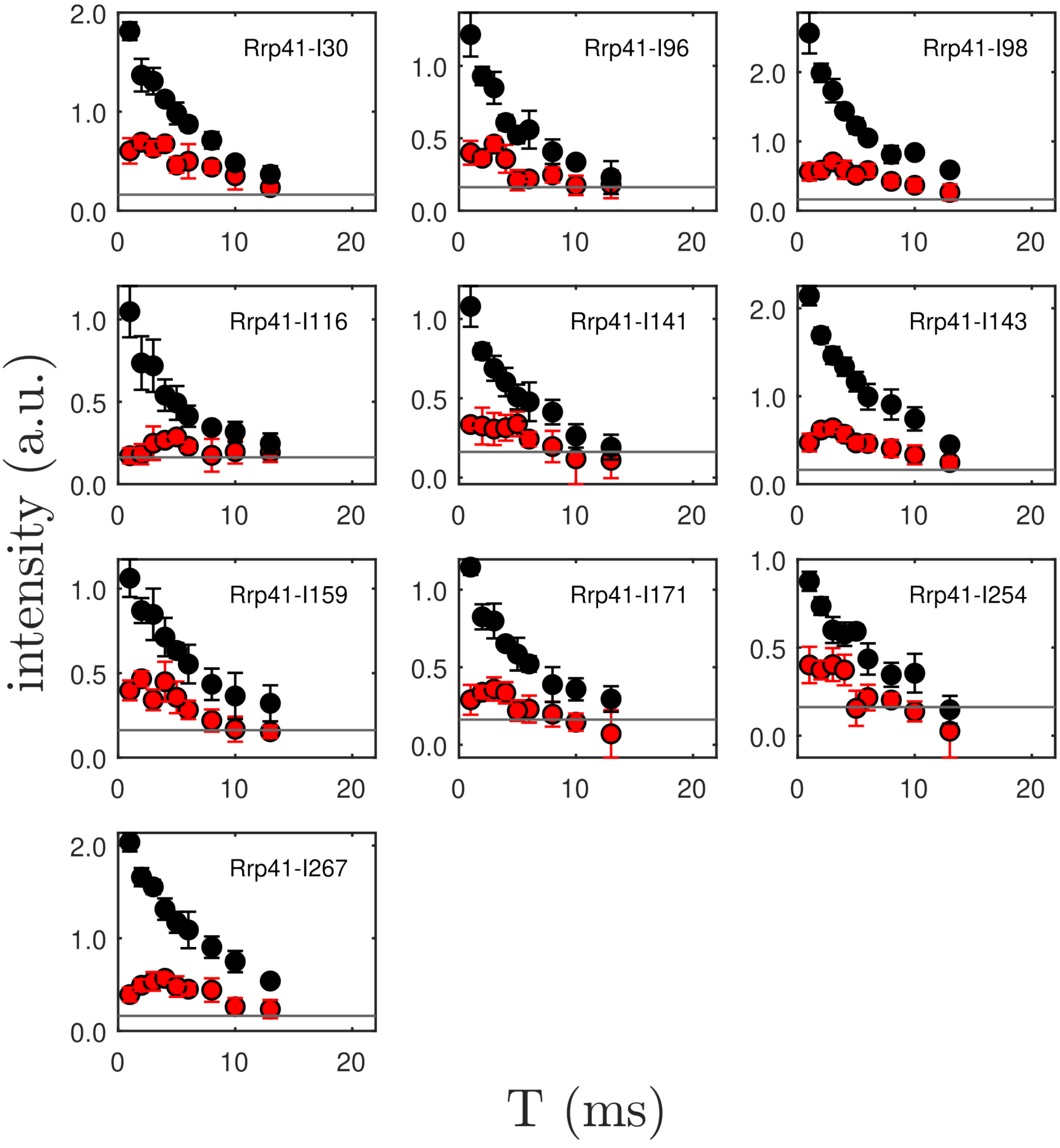
Rrp41-I30
Rrp41-I96
Rrp41-I98
Rrp41-I116
Rrp41-I141
Rrp41-I143
Rrp41-I159
Rrp41-I171
Rrp41-I254
Rrp41-I267
intensity (a.u.)
T (ms)

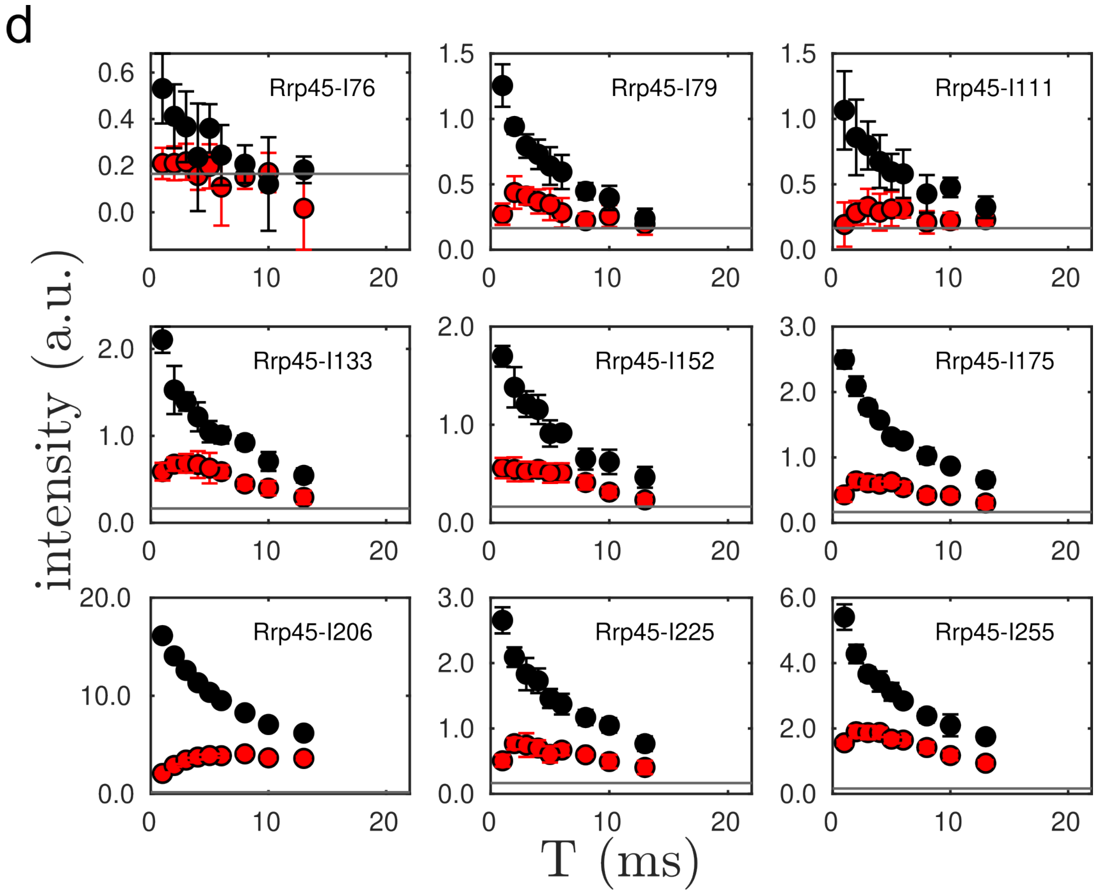


**Figure S4: SQ decay (black) and TQ build-up (red) for Ile-δ1 methyl groups in (a) Csl4, (b) Rrp40, (c) Rrp41 and (d) Rrp45 reconstituted into Exo9 with RNA.** The horizontal grey line indicates the noise level. Data below the noise level was not considered in subsequent analyses. Note that the assignment for residues Csl4-I60/Csl4-I95, Rrp41-I117/Rrp41-I251 and Rrp45-I206 are ambiguous. Csl4-I60 and Csl4-I95 overlap, the assignment for Rrp41-I117 and Rrp41-I251 may be reversed and Rrp45-I206 overlaps with one or more unassigned resonance.

102

103 a

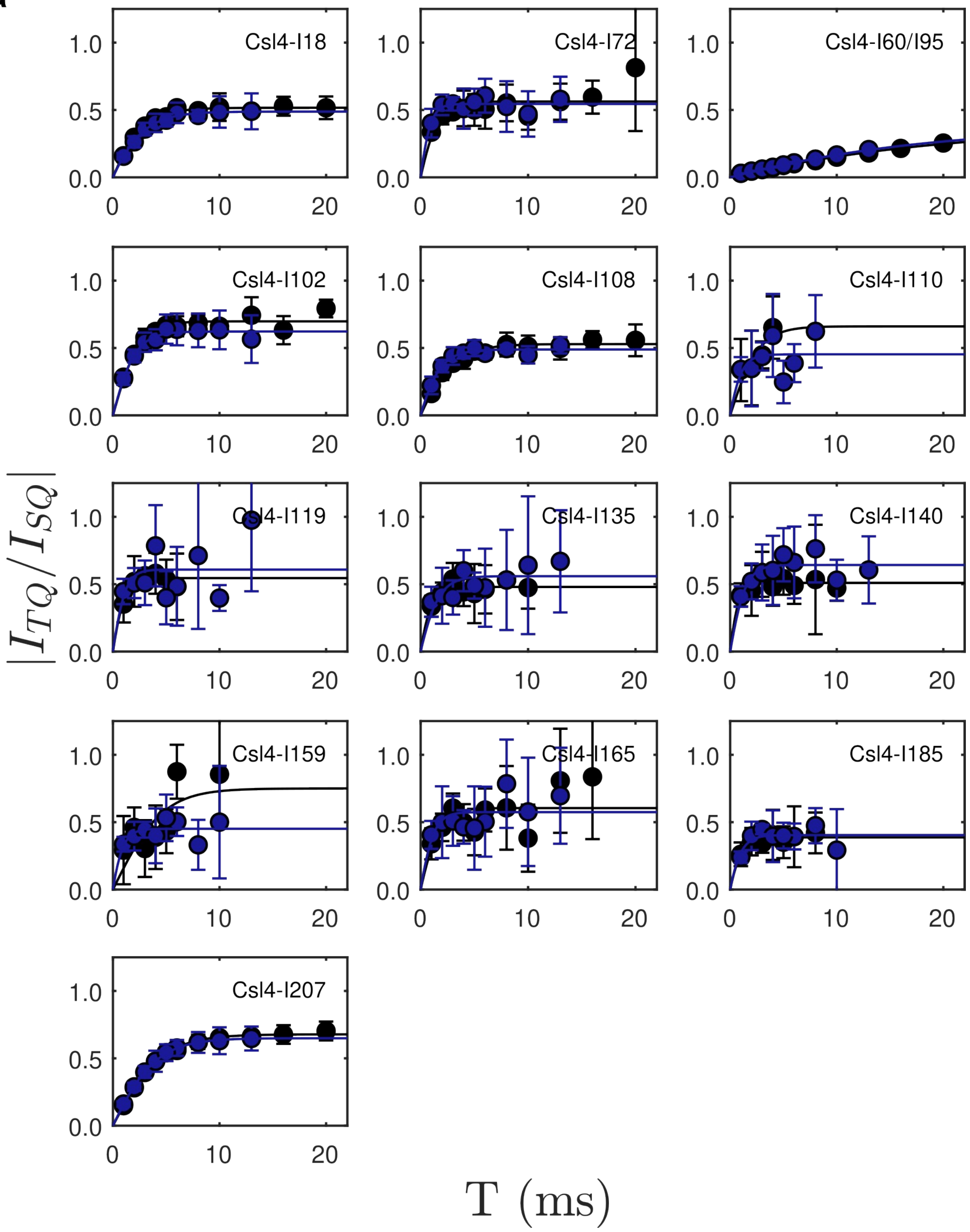

Csl4-I18
Csl4-I72
Csl4-I60/I95
Csl4-I102
Csl4-I108
Csl4-I110
Csl4-I119
Csl4-I135
Csl4-I140
Csl4-I159
Csl4-I165
Csl4-I185
Csl4-I207
1.0
0.5
0.0
0
10
20
$|I_{TQ}/I_{SQ}|$
T (ms)

104

105

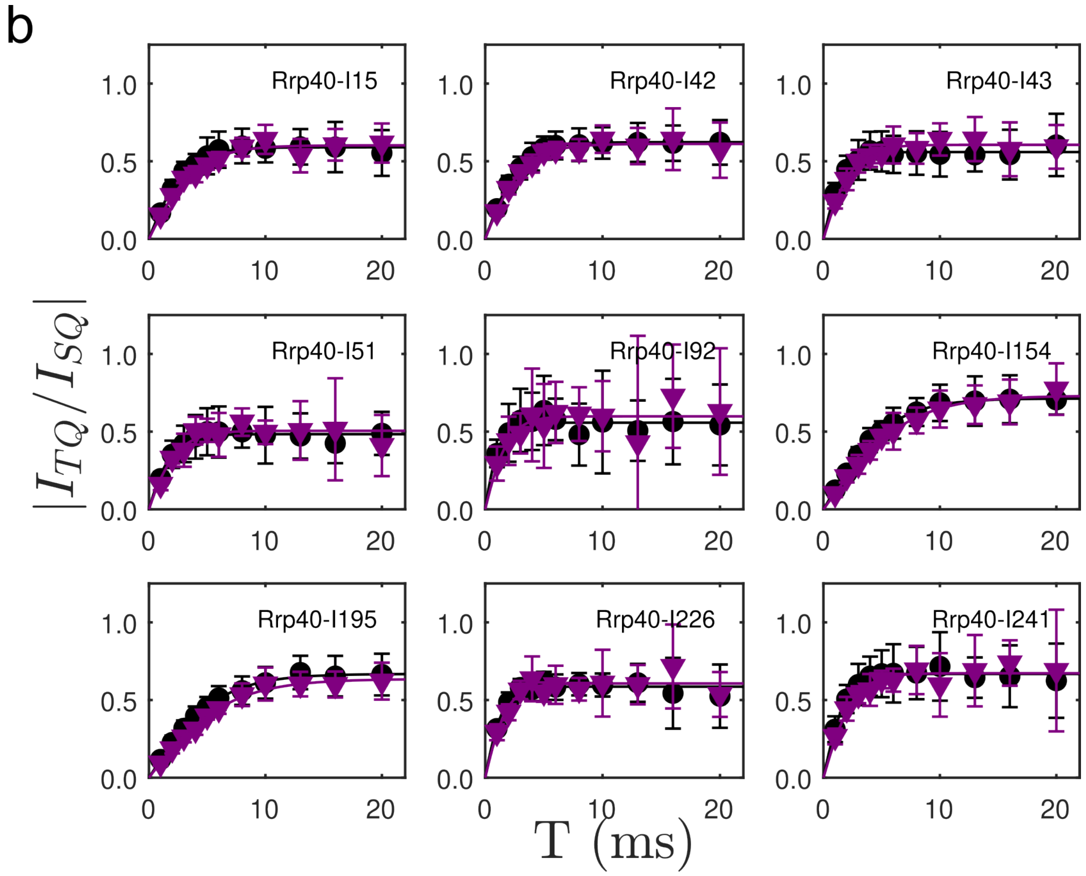

b
Rrp40-I15
Rrp40-I42
Rrp40-I43
Rrp40-I51
Rrp40-I92
Rrp40-I154
Rrp40-I195
Rrp40-I226
Rrp40-I241
$|I_{TQ}/I_{SQ}|$
T (ms)

C

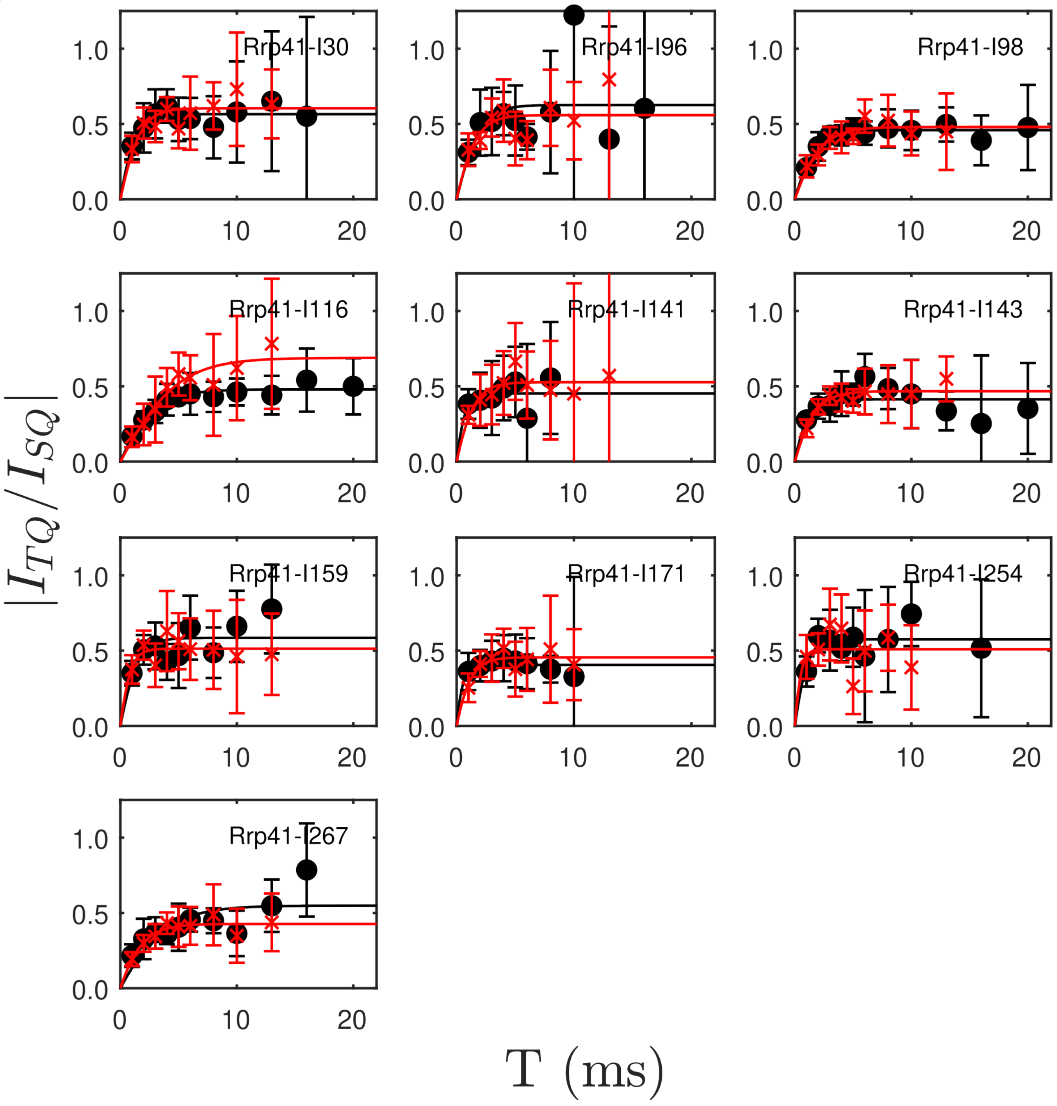

Rrp41-I30
Rrp41-I96
Rrp41-I98
Rrp41-I116
Rrp41-I141
Rrp41-I143
Rrp41-I159
Rrp41-I171
Rrp41-I254
Rrp41-I267
$|I_{TQ}/I_{SQ}|$
T (ms)


107

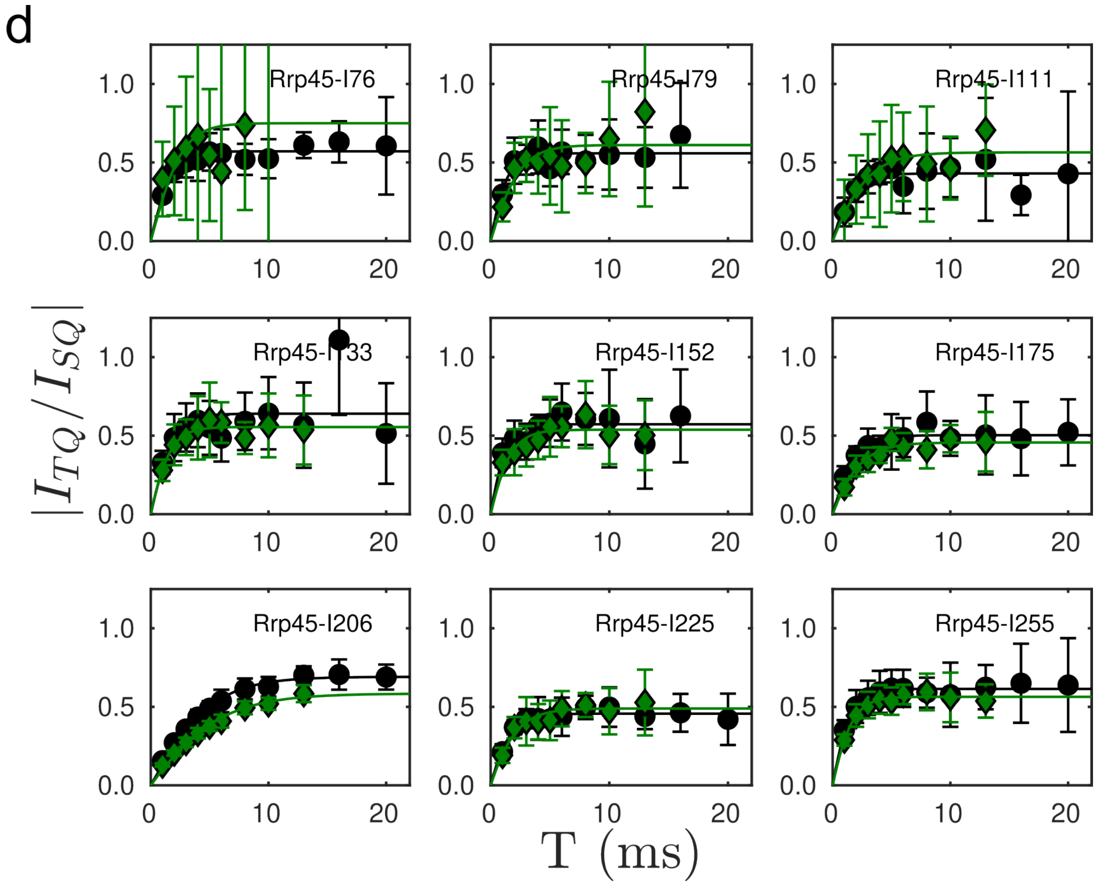


**Figure S5: Order parameter build-up curves for Ile-δ1 methyl groups in (a) Csl4, (b) Rrp40, (c) Rrp41 and (d) Rrp45 reconstituted into Exo9 without (black) and with RNA (Csl4: blue spheres, Rrp40: purple triangles, Rrp41: red crosses, Rrp45: green diamonds).** Note that the assignment for residues Csl4-I60/Csl4-I95, Rrp41-I117/Rrp41-I251 and Rrp45-I206 are ambiguous. Csl4-I60 and Csl4-I95 overlap, the assignment for Rrp41-I117 and Rrp41-I251 may be reversed and Rrp45-I206 overlaps with one or more unassigned resonance.

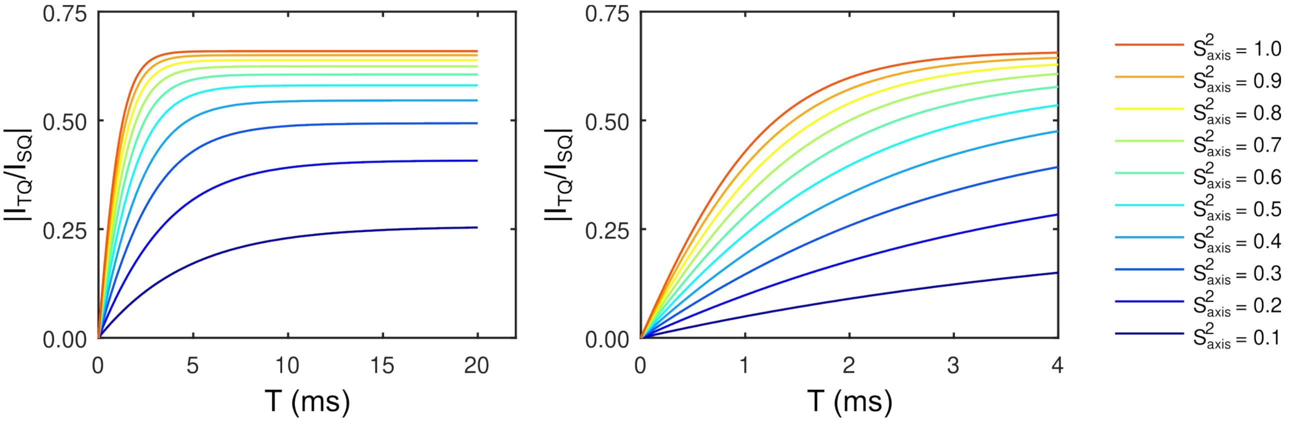


**Figure S6: Simulated build-up curves for order parameters between 0.1 and 1.0 (left) and a zoom into the initial 4 ms regime of the build-up curve (right).** Simulation parameters were: $S_{axis}^2$ = [0.1, 0.2, 0.3, 0.4, 0.5, 0.6, 0.7, 0.8, 0.9, 1.0], $\tau_C$ = 200 ns and $r_H$ = 2.5 Å.

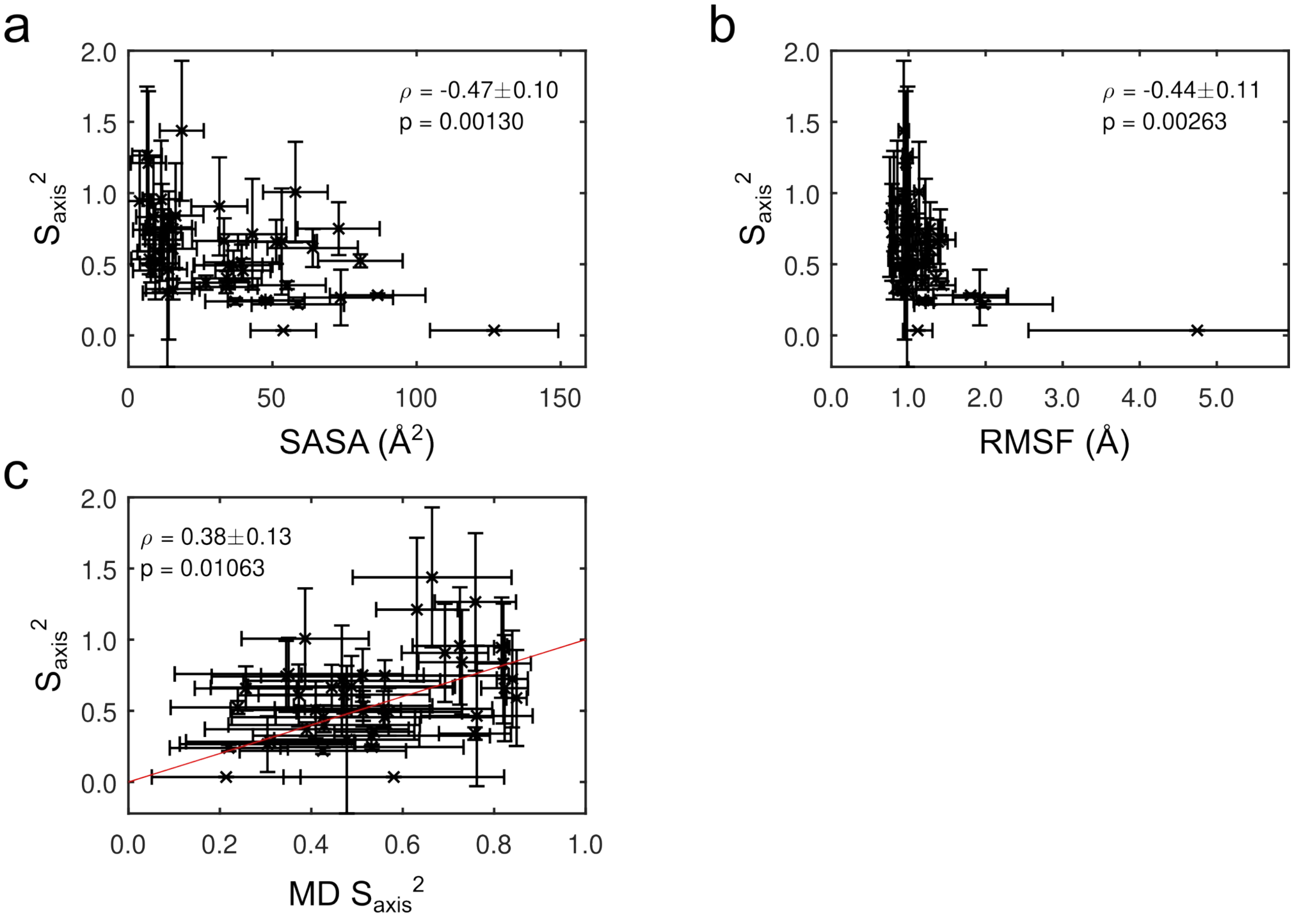


**Figure S7: Correlation between axial methyl order parameters and (a) SASA, (b) RMSF and (c) order parameters calculated from MD simulations.** Spearman's rank correlation coefficient ρ is indicated ±1 SD and *p*-values were evaluated using a two-sided paired-sample *t*-test. The maximum expected ρ, given the measurement uncertainties and distribution of $S_{axis}^2$, is $\rho_{ceil}$ = 0.72±0.07.

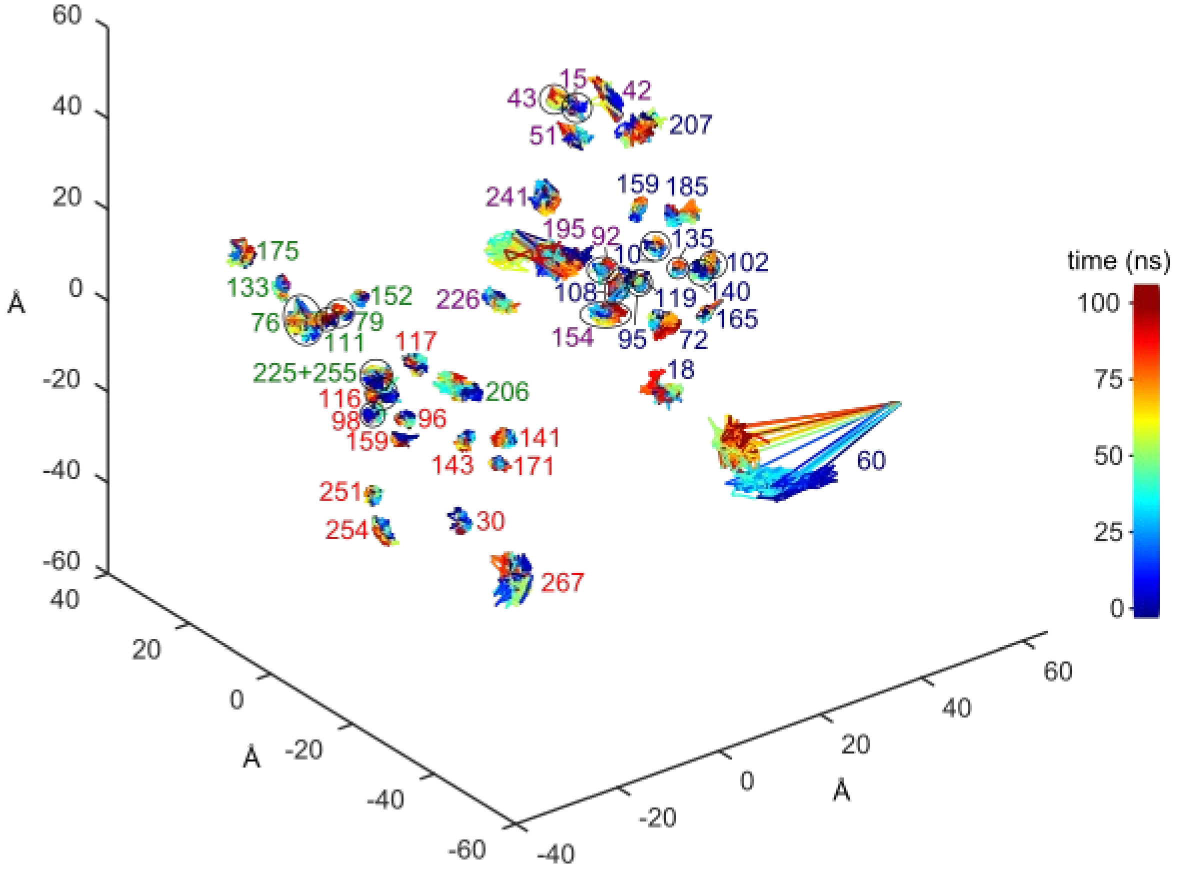


**Figure S8: Space sampled by the Ile-δ1 carbon atoms of Csl4 (residues labeled in blue), Rrp40 (residues labeled in purple), Rrp41 (residues labeled in red) and Rrp45 (residues labeled in green) in Exo9 during a 100 ns MD simulation.** Residues, for which methyl spectra could not be assigned (Rrp45-I168, Rrp45-I233, Rrp45-I236, Rrp40-I225), are not shown.

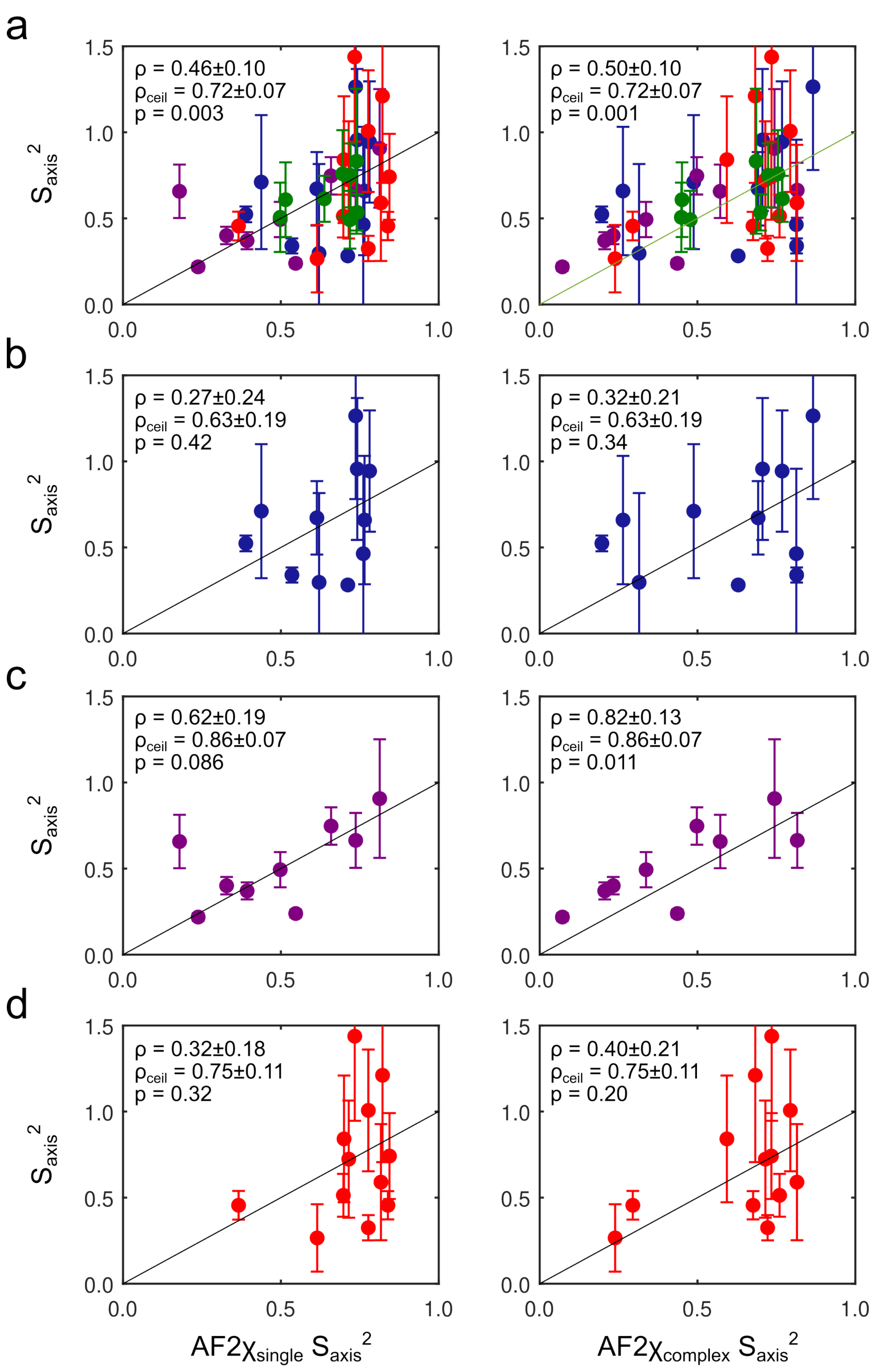
a
ρ = 0.46±0.10
ρceil = 0.72±0.07
p = 0.003
ρ = 0.50±0.10
ρceil = 0.72±0.07
p = 0.001
b
ρ = 0.27±0.24
ρceil = 0.63±0.19
p = 0.42
ρ = 0.32±0.21
ρceil = 0.63±0.19
p = 0.34
c
ρ = 0.62±0.19
ρceil = 0.86±0.07
p = 0.086
ρ = 0.82±0.13
ρceil = 0.86±0.07
p = 0.011
d
ρ = 0.32±0.18
ρceil = 0.75±0.11
p = 0.32
ρ = 0.40±0.21
ρceil = 0.75±0.11
p = 0.20
Saxis2
AF2χsingle Saxis2
AF2χcomplex Saxis2

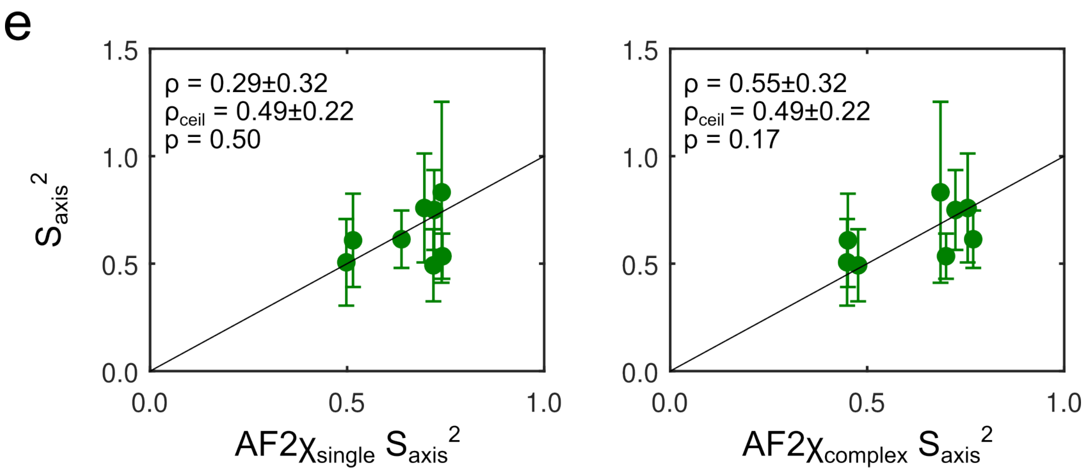


**Figure S9: Correlation between experimental and AF2χ-predicted axial methyl order parameters for (a) all residues, (b) Csl4, (c) Rrp40, (d) Rrp41, and (e) Rrp45 using the 'single' (left) and 'complex' (right) approach as described in the method section.** The black line indicates values for which $S_{axis}^2 = S_{axis}^2{}_{\mathrm{AF}\chi}$. Spearman's rank correlation coefficient ρ is indicated ±1 SD and *p*-values were evaluated using a two-sided paired-sample *t*-test. The maximum expected ρ, given the measurement uncertainties and distribution of $S_{axis}^2$, $\rho_{ceil}$, is indicated in each plot.

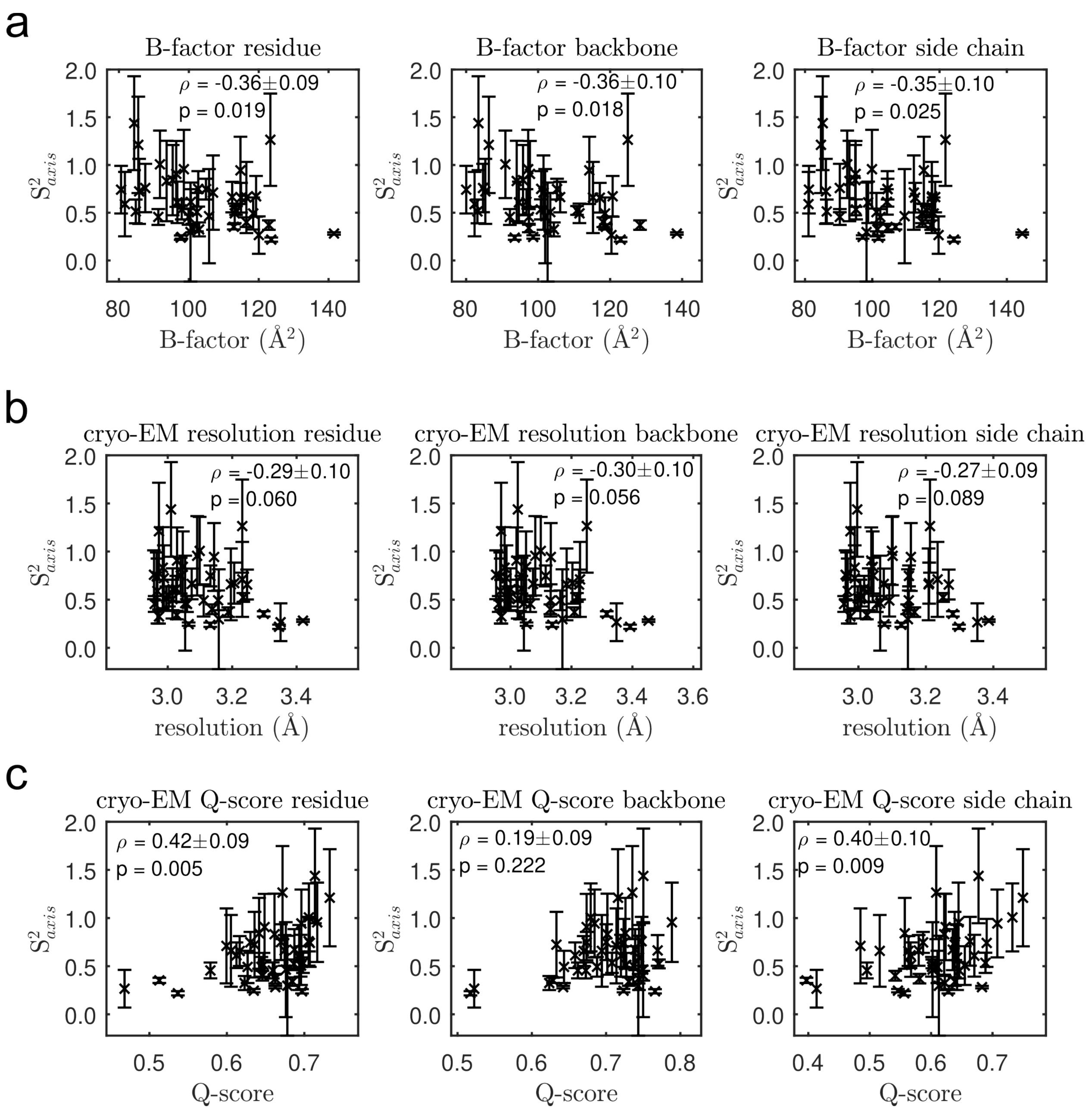


**Figure S10: Correlation between methyl order parameters and various measures of structural resolution. (a)** Correlation between methyl order parameters and X-ray B-factors for the entire Ile residue (left), Ile backbone (center) and Ile side chain (right). **(b)** Correlation between methyl order parameters and cryo-EM resolution of the entire Ile residue (left), Ile backbone (center) and Ile side chain (right). **(c)** Correlation between methyl order parameters and cryo-EM Q-factors of the entire Ile residue (left), Ile backbone (center) and Ile side chain (right). Spearman's rank correlation coefficient ρ is indicated ±1 SD and *p*-values were evaluated using a two-sided paired-sample *t*-test. The maximum expected ρ, given the measurement uncertainties and distribution of $S_{axis}^2$, is $\rho_{ceil}$ = 0.74±0.06.

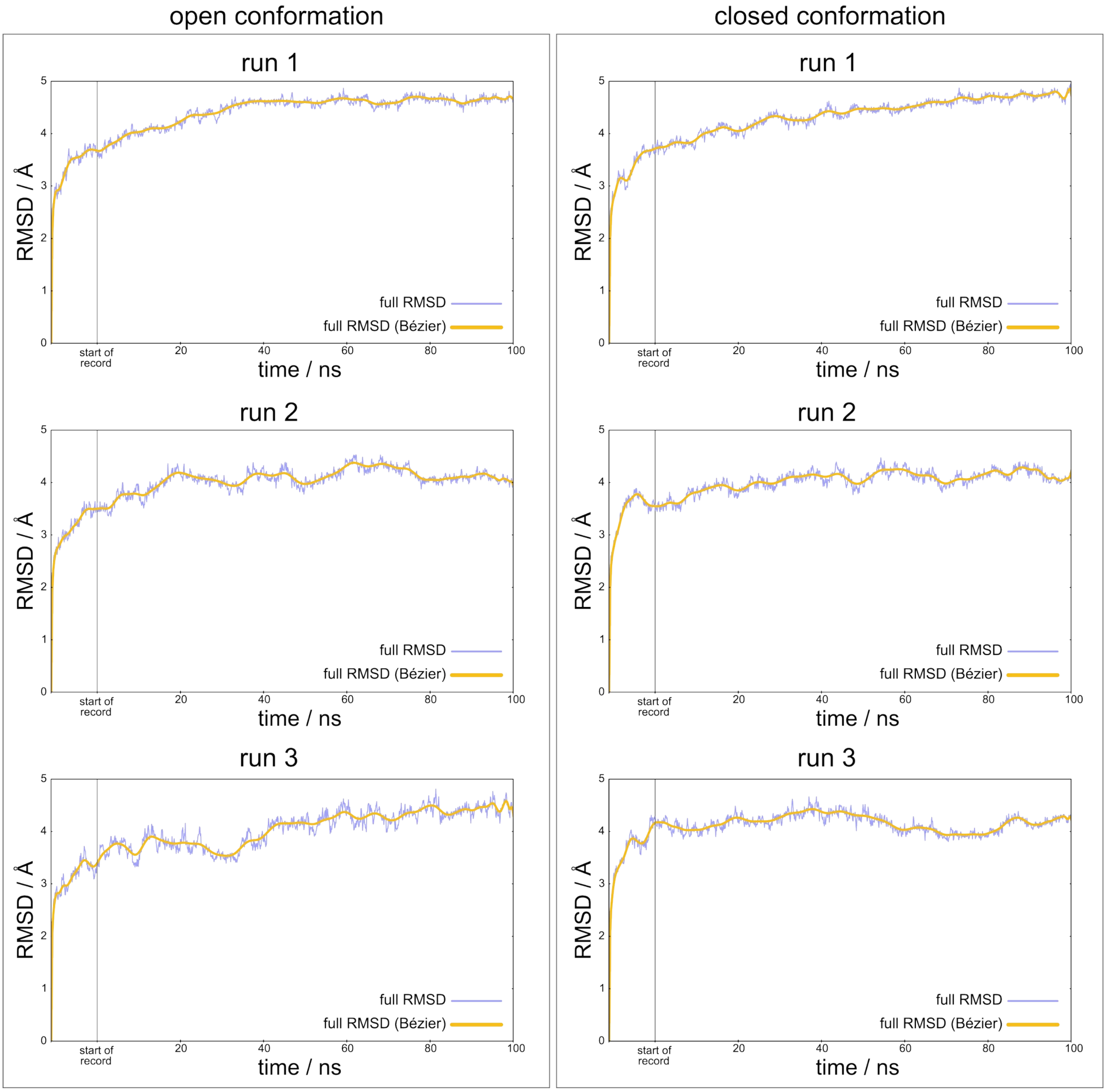


**Figure S11:Structural stability of the nonameric exosome complex during MD simulation.** Time evolution of the Cα root-mean-square deviation (RMSD) for three independent replicas of the exosome complex in its open (left) and closed (right) states. For each trajectory, the vertical gray line marks the transition from the pre-production phase (heating, NVT and NPT ensembles, shown to the left of the line) to the production run (shown to the right). RMSD values were calculated relative to the respective energy-minimized pre-heating starting structure after least-squares fitting of the Cα atoms. In all six trajectories, the RMSD plateaus by the second half of the production run, indicating that the systems had reached energetic equilibrium and validating the use of this portion of the trajectories for subsequent analysis.
